# Privileged representational axes in biological and artificial neural networks

**DOI:** 10.1101/2024.06.20.599957

**Authors:** Meenakshi Khosla, Alex H Williams, Josh McDermott, Nancy Kanwisher

## Abstract

How do neurons code information? Recent work emphasizes properties of population codes, such as their geometry and decodable information, using measures that are blind to the native tunings (or ‘axes’) of neural responses. But might these representational axes matter, with some privileged systematically over others? To find out, we developed methods to test for alignment of neural tuning across brains and deep convolutional neural networks (DCNNs). Across both vision and audition, both brains and DCNNs consistently favored certain axes for representing the natural world. Moreover, the representational axes of DCNNs trained on natural inputs were aligned to those in perceptual cortices, such that axis-sensitive model-brain similarity metrics better differentiated competing models of biological sensory systems. We further show that coding schemes that privilege certain axes can reduce downstream wiring costs and improve generalization. These results motivate a new framework for understanding neural tuning in biological and artificial networks and its computational benefits.

## Introduction

The quest to unravel the neural basis of perception is guided by two opposing paradigms. One view holds that neural representations are best understood at the level of an entire population of neurons [1–6]. Under this view, the manner in which individual neurons realize neural representations is deemed inconsequential, and the coordinate axes of the representation thus hold no relevance: a representational system, denoted as A, is treated equivalently to another system, B, derived by rotating the axis of the original representation, A. This view is supported by the argument that changing the coordinate axes preserves both linearly decodable information and representational geometry (the relative distances among patterns of response in representational space). An alternate view contends that the instantiation of a representation at the level of individual neurons matters [1, 7–9], and System A and System B could differ importantly from each other (see Fig. 1).

**Fig. 1.**
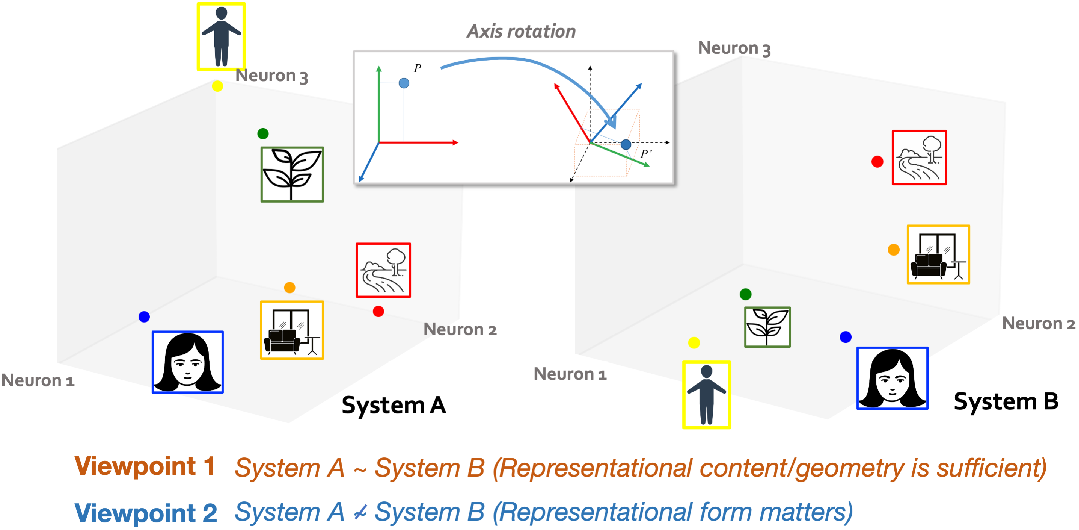
Cartoon schematic presenting two representational frameworks in three dimensions. Each stimulus elicits a unique response pattern in the three units. In System A, the cardinal axes correspond to individual neurons, where neuron 1, 2, and 3 exhibit selective responses to faces, scenes, and bodies, respectively. In System B, we depict the representation achieved by arbitrarily rotating the coordinate axes of System A. In this configuration, the units do not exhibit clear tuning to any of the categories.

How can one resolve the apparent conflict between these two viewpoints, and decide which framework is more suitable for understanding the neural code? Two questions must be answered. One is empirical: Do brain regions have a preferred way of implementing representations at the level of individual neurons? The second question is theoretical: Why might the implementation of a representation at the neuron level matter, that is, what are the consequences of choosing a particular representational axis?

Decades of experimental investigations have provided loose evidence bearing on the empirical question. Individual neurons, and sometimes whole brain regions, have been reported to encode complex, meaningful and identifiable features of our external world such as the presence of a navigational boundary, face, or speech sound [10–15]. These and many other selective response profiles are consistently found across individuals and sometimes even across species [12, 16–25], raising the possibility that some brain systems might favor certain representational coordinate systems over others. However, this body of evidence remains predominantly qualitative, and leaves open the possibility that beyond a few specific tuning functions, axes could mostly differ across brains. Thus, an important question remains unresolved: Are neural representational axes arbitrary, or do neural populations in fact code for information in more systematic ways, such that responses of individual neurons might be aligned across brains?

Many data analysis practices in modern neuroscience implicitly assume that representational axes are irrelevant, and are therefore poorly suited to address this question. For example, linear decoding models are insensitive to arbitrary rotations of the neural representational axes: imposing a rotation on the representations will simply transform the optimal decoding weight matrix and lead to no change in predictive accuracy. Similar assumptions are built into the most popular methods for testing the alignment of representations across brains and artificial networks, including representational similarity analysis (RSA) [26, 27], canonical correlation analysis (CCA) [28], centered kernel alignment (CKA) [29, 30], Procrustes shape distance [30] and linear predictivity scores [31]. Thus, while such comparative analyses of biological and artificial neural networks have been instrumental in explaining the “*content*” of neural codes (i.e. what information is linearly accessible from a representation and its representational geometry), they fail to explain the “*form*” of neural codes (i.e. how individual neurons encode information) due to their invariance to rotations of response patterns. Moreover, these analytical tools implicitly assume a down-stream readout that has access to the entire neural population and is capable of computing linear transforms. However, the accuracy of this assumption is yet to be determined.

Here, we show that the view of neural and artificial coding that disregards the significance of representational axes is incomplete at best, as it cannot account for a striking empirical fact: certain axes of representation are “privileged”, that is, systematically aligned across brains and across DCNNs trained to process natural visual and auditory inputs. To assess this alignment, we introduce measures for quantifying alignment between representations that overcome the constraints imposed by existing rotation-insensitive comparison metrics. These axis-dependent alignment measures reveal alignment across conspecifics’ perceptual cortices, across different DCNNs and between perceptual cortices and respective DCNNs in their native axes. Remarkably, in most cases, this alignment approaches the maximal alignment achievable through any axis rotation. These findings held in both vision and audition, and our preliminary analyses suggest they also extend to more cognitive representations including language. Further, our alignment measures, which map individual neurons or voxels to individual network units, establish a more stringent match between brains and artificial networks, and better discriminates model fit to the brain compared to currently standard methods like regression-based model prediction. Finally, we demonstrate that the axes privileged in these coding schemes convey computational benefits in terms of downstream wiring costs, sparse coding and the capacity for few-shot generalization under biologically realistic constraints.

## Results

We present a method for assessing whether and to what extent a representational system has privileged axes, and then use this method to ask the following questions: 1) Do sensory cortices (both visual and auditory) have privileged representational axes?, 2) Do deep convolutional neural network models trained on natural visual and auditory input also have privileged axes? 3) Do biological and artificial neural networks share the same representational axes?, 4) Are any of these aligned basis directions interpretable?, 5) Does brain-model axis alignment improve our ability to choose between competing models of biological computations (e.g. DCNNs vs. transformers)?, 6) Are there computational advantages to the axes that are privileged? and 7) Are privileged axes also found in higher-order cognitive domains?

### A formal framework for assessing privileged axes

Consider a representation 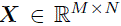 formed by collecting the responses of *N* neurons to *M* stimuli. The activation space corresponds to ℝ^*N*^. The native basis 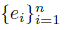 (with *ei* defined such that its *ith* entry equals 1 and all others equal 0) is but one of an infinite number of possible bases for ℝ^*N*^. Any geometry-preserving change of basis can be constructed by sampling a rotation matrix (i.e. a special orthogonal matrix) 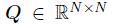 and applying the corresponding linear transformation to ***X***, yielding the transformed representation ***XQ***. Critically, this change of basis preserves the information *content* of a representation (i.e., what can be linearly decoded) as well its *geometry* (i.e. distances between points). This raises the question: are the native basis and the axes it defines arbitrary?

Here, we address this question using a formal framework for assessing *whether* and *to what extent* a representational system (e.g. a brain area or DCNNs) has privileged representational axes. We test for such axes by examining multiple instances of a given representational system (e.g. different individuals of a species, or different instantiations of artificial neural networks) and asking whether there is any consistent relationship between their axes.

In what follows, we first introduce axis-dependent measures of representational alignment. We then use them to measure the alignment between instances of brains or artificial neural networks and assess how the alignment changes under axis perturbations. If the pair of representation instances are consistently more aligned in their native basis than in the perturbed bases, this provides evidence that the representational scheme has privileged axes. We then contrast these values with the maximal alignment achievable under all possible bases. If the alignment in the native basis approaches the maximal alignment under all possible bases, that provides further support for privileged axes.

We note that by “privileged axis”, we refer exclusively to a consistent inclination towards certain bases as evident across different system instances. Privileged axes need not be interpretable, or to have other distinguishing properties. In later sections, however, we consider the properties of the privileged axes and whether those might explain why they are privileged.

#### Quantifying representational alignment

We introduce two complementary measures of axis-alignment. The first measure is an intuitive quantification of direct pairwise correspondence between tuning functions in different representations. The second measure is symmetric and benefits from the desirable properties of metric spaces in its theoretical framework.

##### Pairwise matching score

Our first measure is a direct measure of similarity between units in different representations. Here, a unit in a biological representation corresponds to the smallest unit of measurement in brain recordings (a voxel in fMRI data, a single neuron in neurophysiology data), whereas a unit in artificial networks corresponds to a single “filter” in the convolutional layer of a DCNN with its response averaged over all spatial locations.

Consider two matrices 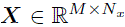 and 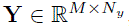, which represent the activations of *Nx* and *Ny* units respectively, for M inputs (e.g. stimuli). The proposed measure proceeds in two stages:

- In the first stage, we find the optimal assignment matrix 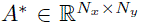 with entries 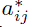 that maps each unit in ***X*** to a single unit in ***Y*** by optimizing a correlation-based objective. The optimization problem is defined as follows,

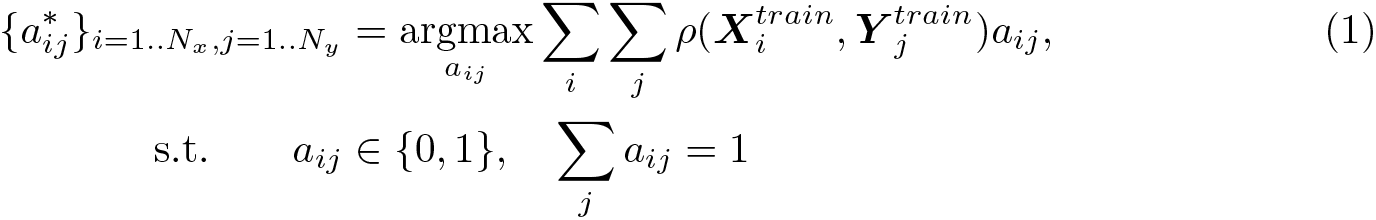 Intuitively, the optimization procedure aims to find, for each unit in ***X***, the unit in ***Y*** that exhibits the maximum response correlation with that unit in ***X*** over a training stimulus set. The constraints on *aij* ensure that each unit in ***X*** is matched to exactly one unit in ***Y*** (although each unit in ***Y*** can have multiple units in ***X*** mapped to it). As a result, some units in ***Y*** may remain unmatched after the assignment.
- In the second stage, using the unit matches obtained in the first stage, we compute the correlation between each unit in ***X*** and its corresponding matched unit in ***Y*** on held-out data and then average these values across all matched pairs to derive the final alignment metric

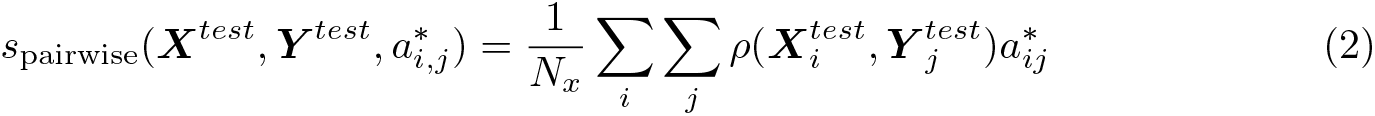

Note that this measure is asymmetric by design. The choice to allow repetitions with a bipartite semi-matching assignment (i.e. allowing a single unit in ***Y*** to be matched to multiple units in ***X***) [32] instead of a strict bipartite matching is motivated by potential variations in the number of neurons/units across different systems.

##### Soft matching score

The pairwise matching score just described is intuitive but is inherently asymmetric: we can obtain different alignment values when aligning units from ***X*** to ***Y***, compared to the reverse. Due to the asymmetry, it could also yield counter-intuitive outcomes like deeming a representation ***X*** with many units as somewhat aligned to any representation that is mapped to it [33].

Motivated by these limitations, we employ another novel axis-dependent representational similarity measure, rooted in optimal transport theory, to quantify the similarity in the tuning curves of units across two representations. Intuitively, this alignment measure is a generalization of the permutation-based similarity measure proposed in Williams et al. [30] that optimizes over the set of neuron permutations to align two representations. Here, this generalization employs a “soft-permutation” approach instead to match units across two systems with different numbers of neurons (see [33] and Methods for more background on this similarity measure). Through empirical simulations, we have shown that both the pairwise matching and soft-matching alignment measures effectively capture similarity in the “form” of representations, above and beyond “content” comparisons (see Supplementary Fig. S1).

Our main findings in this paper hold for both the pairwise matching and soft matching alignment metrics. In the main text, we show results derived using the pairwise matching score, while results based on the soft matching score are provided in the Supplementary Materials (see Supplementary Figures S4, S5 and S6).

#### Probing alignment changes with axis perturbations

To test whether brains and models have privileged axes, we ask whether the alignment between instances is greater than would be expected from an arbitrary axes. We draw a random rotation matrix ***Q*** uniformly from the special orthogonal group *SO*(*n*). We also consider smaller rotations (closer to the identity matrix ***I***) along the SO(n) manifold by computing fractional powers of the sampled rotation matrix, ***Q****α*(0 *< α <* 1). For each pair of representation instances, we rotate the axes of one instance along this SO(n) manifold and quantify how the alignment between instances changes with the amount of rotation (*α*). We note that each rotated representation has exactly the same linearly decodable information and population geometry and differs only in the tuning of its coordinate axes.

#### Deriving maximal alignment across all axis rotations

The aforementioned methodology tests whether the alignment between axes in different instances of the system is higher in their native basis compared to a uniformly drawn basis. But it leaves open the possibility that some rotation matrix ***Q***∗ in the SO(n) manifold yields a higher alignment than the native basis. We use the Lie exponential and perform gradient-based optimization on the SO(n) matrix manifold [34] to find ***Q***∗ and compute this maximal alignment for both alignment measures *s* ∈ {*s*softmatch*, s*pairwise}.

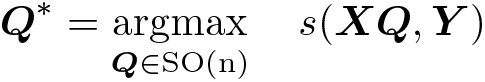

See Methods for further details. To evaluate whether the native basis alignment *s*(***X****, **Y***) approaches the maximal basis alignment *s*(***XQ***∗ *, **Y***), we express the basis alignment as a proportion of the maximal possible value.

We validate the measures of alignment with empirical simulations that show the framework to be effective in detecting the presence or absence of privileged axes in various representational systems (see Supplementary Figures S2 and S3).

### Sensory cortices have privileged axes

We first tested whether perceptual cortices exhibit privileged axes for the representation of natural stimuli. To answer this question for the visual system, we leveraged the Natural Scenes Dataset (NSD, human fMRI) [35] and the ManyMonkeys dataset (macaque neurophysiology) [36] to assess how axis-dependent alignment between conspecifics’ neural responses in the human ventral visual cortex and macaque IT changes with axis rotation respectively. To address the same question in the auditory system, we analyzed a dataset of fMRI responses to natural sounds (measured in the auditory cortex of multiple subjects [37]).

We found that visual representations (both human fMRI and monkey neurophysiology) exhibit near-maximal alignment between individuals (values approaching 100% in Figs. 2B-C), and this alignment decreases gradually with increasing rotation, indicating a non-random relationship between the two bases (Fig. 2B-C). fMRI responses to natural sounds showed a similar pattern, providing evidence that the auditory cortex also has privileged axes for representing auditory stimuli (Fig. 2D). In each of the three datasets, we find that the majority of neurons/voxels lose this direct alignment with neurons/voxels in conspecifics upon axis rotation (Fig. 2B-D, bottom). We also conducted this alignment analysis by sampling rotation matrices through a different method —specifically, as a composition of plane rotations. This approach provides finer control over the degree of rotation and the number of plane (2D) rotations involved. Our findings indicate that even minor perturbations to the native axes result in significant loss in alignment between conspecifics (see Methods and Supplementary Fig. S9). Our results thus show that these biological representational systems have a privileged set of axes, challenging widespread assumptions about their arbitrariness.

**Fig. 2.**
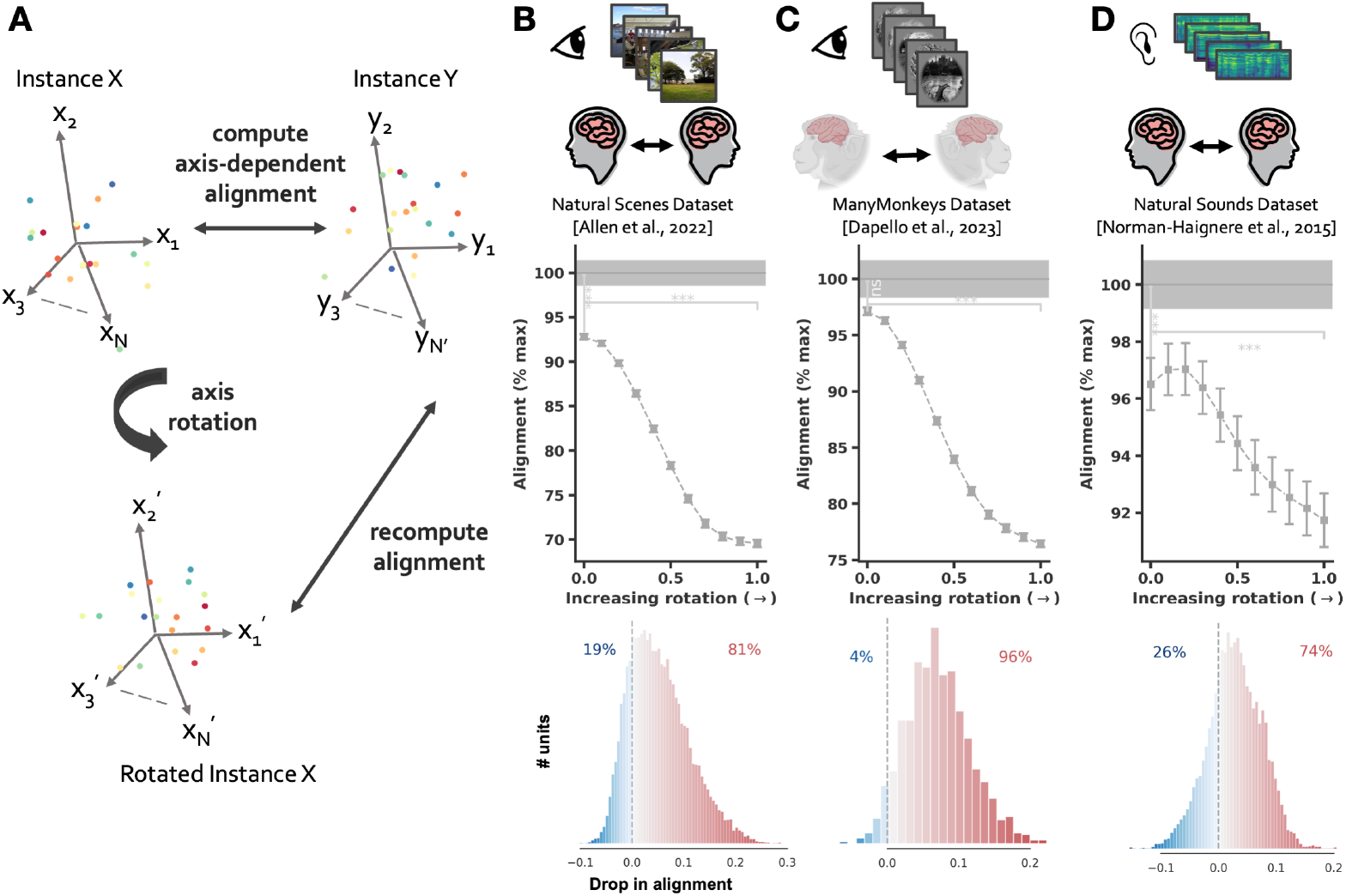
Brain-to-brain alignment. (**A**) Cartoon schematic presenting the framework used to probe the prevalence of privileged axes in sensory cortical representations. Colored dots represent individual stimuli and axes *x*1*, …, xN* and *y*1*, …, yN* represent responses of individual voxels or neurons. Neural representations in different conspecifics are compared with axis-alignment measures sensitive to single-neuron tuning. For each representation pair, we change the axes of one representation along a smooth SO(n) manifold and quantify how the pairwise matching alignment changes with the amount of axis rotation. (**B**), (**C**) and (**D**) depict alignment between conspecifics neural responses in human ventral visual cortex, macaque IT and human auditory cortex. This alignment is measured using the pairwise matching score and examined as a function of axis rotation (*α*) where *α* = 0 indicates no rotation and *α* = 1 indicates full rotation using a rotation matrix drawn uniformly from SO(n). Alignment is expressed as a proportion of the maximum possible value obtainable across all axis rotations, determined using the ‘Optimal Axis Alignment’ procedure described in the text. Histograms depict the distribution of the difference in pairwise matching scores between the native and rotated basis across all neurons/voxels with the bars colorcoded based on difference values. Error bars are SEM over multiple splits of the data. Significance tests compare the alignment values at no rotation (*α* = 0) with those at full rotation (*α* = 1), as well as with the optimal alignment values observed across all rotation angles.

### Deep convolutional networks have privileged axes

To further understand the privileged axes of biological sensory systems, we next asked whether and when artificial neural networks trained on natural stimuli also develop privileged axes.

In the domain of vision, we selected four standard deep convolutional networks, each trained on ImageNet [38], that span different architectures and learning objectives. These include: (i) ResNet50 with supervised training, henceforth called ResNet50 (sup.) [39], (ii) AlexNet with supervised training, denoted as AlexNet (sup.) [40], (iii) ResNet50 trained in a self-supervised fashion with a contrastive learning objective, specifically momentum contrast [41], denoted as ResNet50 (self-sup.) and (iv) ResNet50 with adversarial training, henceforth ResNet50 (robust) [42]. All these models perform competitively on large-scale object categorization. We restrict our analysis to later layers of DCNNs as these are known to best model the high-level visual cortex.

Even when comparing models with different architectures (e.g. ResNet vs. AlexNet) and learning objectives (e.g. supervised vs. self-supervised), we find that DCNNs are almost maximally aligned with each other in their native basis (Fig. 3A), indicating that they contain privileged axes. Importantly, untrained networks do not exhibit this phenomenon (see Supplementary Fig. S7).

**Fig. 3.**
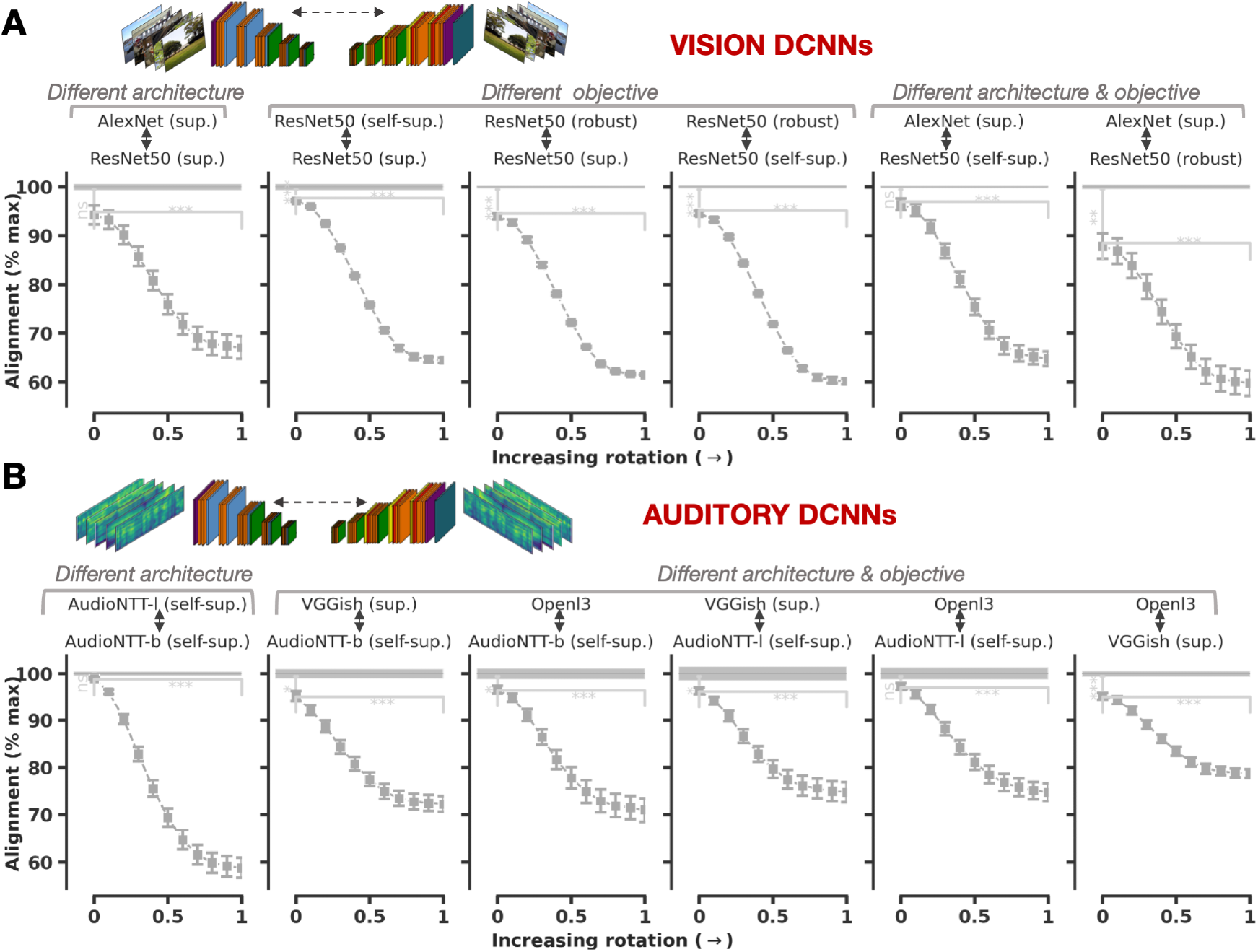
Model-to-Model alignment. (**A**) Alignment between different visual deep convolutional neural networks trained with different architectures and learning objectives. This alignment is measured using the pairwise matching score and examined as a function of axis rotation (*α*) where *α* = 0 indicates no rotation and *α* = 1 indicates full rotation using a rotation matrix drawn uniformly from SO(n). Alignment is expressed as a proportion of the maximum possible value obtainable across all axis rotations, determined using the ‘Optimal Axis Alignment’ procedure described in the text. Alignment computed from activations to the 1000-image NSD stimulus set used for computing inter-subject alignment. (**B**) Same as A, but for auditory neural networks, and computed from activations to the 165 natural sounds used for auditory inter-subject alignment analysis. In each case, we gradually rotate the axis in one representation while computing its alignment with the other representation in its native basis. Error bars are SEM over multiple splits of the data. Significance tests compare the alignment values at no rotation (*α* = 0) with those at full rotation (*α* = 1), as well as with the optimal alignment values observed across all rotation angles.

We performed similar comparisons in the auditory domain, employing four different models, all trained on the AudioSet corpus [43]. These include: (i) a VGG-inspired architecture with supervised training, henceforth called VGGish (sup.) [44], (ii) a convolutional architecture with a high-dimensional representation embedding (2048), trained with contrastive learning, labeled here as AudioNTT-l (self-sup.) [45], (iii) a more compact network akin to (ii) but with a reduced representation embedding dimensionality (512), denoted as AudioNTT-b (self-sup.) [45], and (iv) the *L*3-net audio model, trained with self-supervision on an audio-visual correspondence pretext task, denoted as Openl3 [46]. Irrespective of the architecture and learning objectives, auditory DCNNs after training again converge onto privileged axes (Fig. 3B).

Why do different networks have similar bases? The dominant view from prior work is that the basis is an arbitrary and inconsequential aspect of network representations. This idea stems from the notion that any representation within a neural network is equivalent up to a full-rank matrix multiplication because the successive weight matrix can counterbalance any such multiplication, rendering the initial choice of basis irrelevant [29, 47]. However, this degeneracy is altered by the introduction of non-linearities within deep network architectures. Specifically, consider the set of invertible linear transformations that commute with the non-linear operators used in DCNNs, such as Rectified Linear Units (ReLUs). This set does not encompass the entire space of invertible matrices, but rather is constrained to a limited subset within that space. Non-linearities might thus act as symmetry-breaking mechanisms, potentially predisposing networks to favor certain activation bases.

Concretely, consider the representation of a hidden layer within a deep neural network that comprises an activation basis in R*n*. While R*n* offers an infinite number of alternative bases, our objective is to identify those that could in principle be feasibly learned within the hidden layer representations. Mathematically, let 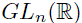 represent the set of all invertible linear transformations in dimension *n* on a vector space over a field of real numbers R. 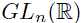 thus consists of all *n* × *n* matrices with real entries and non-zero determinants. Now, for ReLU nonlinearities, consider the subset 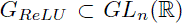 whose action on the network activations prior to ReLU has an equivalent transformation in 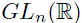 after the pointwise application of this nonlinearity.

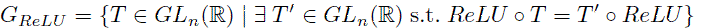

For linear transformations in deep network layers, the set of invertible linear transformations which when applied before the function allow for an equivalent transformation after their application, spans the entirety of 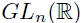. Thus, a completely linear network should not have privileged axes. However, *G_ReLU_* does not encompass 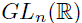, but is substantially restricted to the set of matrices *PD*, where 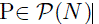 is a permutation matrix and D is a diagonal matrix with positive entries. The integration of ReLUs into the network thus disrupts the existing symmetry and can endow the network with a preferred axis.

### DCNNs and sensory cortices have aligned privileged axes

The presence of privileged axes in perceptual cortices and DCNNs raises the question of whether sensory areas and DCNNs have the same privileged axes. To address this question, we measured the axis-dependent alignment between brain representations and DCNNs, and examined how it changes with increased axis rotations. For the visual domain, we incorporated data not only from the ManyMonkeys dataset and the Human NSD but also from a novel dataset capturing responses in macaque IT to images from the human NSD. We conducted analogous experiments with auditory models and human auditory cortex responses, measured using both fMRI and intracranial recordings (ECoG or electrocorticography) [48].

Across all brain-model pairs in both vision and audition, alignment was high without rotation (often, approaching the maximal value) and decreased by as much as 20 35% as DCNN axes were rotated (Fig. 4A-E). This result suggests that sensory cortices and DCNNs tend to exhibit the same axes.

Are the tuning functions that are most aligned between conspecifics also the ones most aligned between brains and DCNNs? To find out, we asked whether the capacity of a neuron’s or voxel’s tuning function to be accurately represented by that of a single neuron or voxel in a conspecific can predict its peak alignment with the tuning function of a single unit in a DCNN (Fig. 4F). Across neurons/voxels, we found a significant correlation between the two, even after adjusting for the varying reliability of response measurements in different neurons/voxels. This result indicates that the neurons or voxels which exhibit high one-to-one alignment across conspecifics also tend to be more one-to-one aligned with DCNNs.

**Fig. 4.**
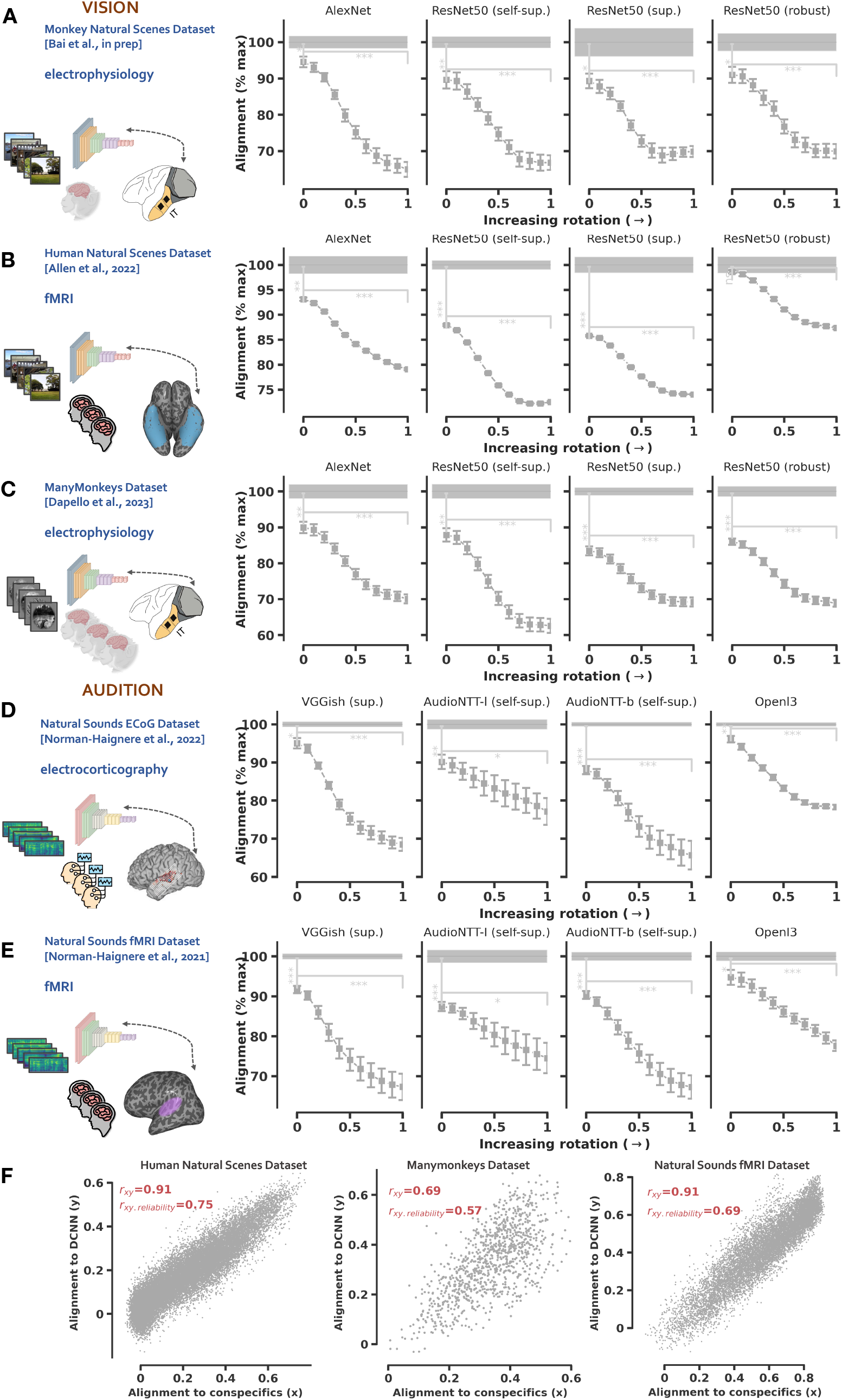
Model-to-brain alignment. (**A-C**) Alignment of various visual deep convolutional neural networks with biological neural representations in the macaque IT and human ventral visual stream. This alignment is measured using the pairwise matching score and examined as a function of axis rotation (*α*) where *α* = 0 indicates no rotation and *α* = 1 indicates full rotation using a rotation matrix drawn uniformly from SO(n). Alignment is expressed as a proportion of the maximum possible value obtainable across all axis rotations, determined using the ‘Optimal Axis Alignment’ procedure described in the text. (**D-E**) Alignment of auditory DCNNs with neural representations in the human auditory cortex, again assessed as a function of DCNN axis rotation. (**F**) illustrates the one-to-one alignment of each neuron/voxel with neurons/voxels in other conspecifics as a function of its alignment with units in a DCNN (self-supervised vision/auditory networks). The inset reports the raw correlation between these variables and the partial correlation after excluding the effect of different noise ceilings of various neurons/voxels. Error bars are SEM over multiple splits of the data. Significance tests compare the alignment values at no rotation (*α* = 0) with those at full rotation (*α* = 1), as well as with the optimal alignment values observed across all rotation angles.

We emphasize that standard approaches for comparing artificial and biological neural representations (linear encoding, RSA or CKA) conceal this similarity in *representational form* between DCNNs and sensory regions. Assessing similarity in form requires measures of axis alignment such as those introduced here.

Why do biological and machine systems converge on similar representational axes? Importantly, untrained networks do not show axis alignment with neural representations (see Supplementary Fig. S8). This result suggests that optimization constrained by natural input statistics and the architectural primitives in DCNNs (e.g. ReLUs) might help explain why brains favor certain coordinate systems over others.

### Privileged axes are category-selective

In neuroscience, a wealth of evidence has revealed the presence of neurons tuned to meaningful features of our external environment. For example, in the macaque inferotemporal cortex, many single neurons selectively respond to images of faces [12]. In humans, both neuroimaging studies and intracranial recordings within the visual cortex have identified neural populations tuned to images of faces, scenes, bodies, visually presented letter sequences or words, and food [13, 16–19]. Distinct neural populations within the auditory cortex analogously show selectivity for music, speech, and song [37, 48]. It has remained unclear why such category-selective tuning is so consistently observed across neural recordings in distinct individuals. The findings in the previous section demonstrating aligned axes between DCNNs and sensory cortices raise the possibility that DCNNs also possess analogous tuning functions.

To explore this issue, we employed a data-driven, hypothesis-agnostic factorization approach to discover the dominant components underlying DCNN representations. The rationale was to reduce the representational dimensionality to a tractable number for interpretability analyses while minimizing biases towards predetermined categories. Models trained with supervision might plausibly acquire category selectivity via training labels that correspond to such categories. Accordingly, we instead analyzed self-supervised models which do not receive any explicit supervision in the form of semantic labels, as these seemed the strongest test of the presence of selectivity related to particular categories.

We first aggregated the responses of all *n* units in the self-supervised visual DCNN (trained with momentum contrast) to *m* natural stimuli from the NSD, resulting in a data matrix ***D*** ∈ R*m*×*n*. We employed Bayesian Non-Negative Matrix Factorization method previously applied to neural data [19] to factorize the DCNN responses to natural stimuli into two lower-dimensional matrices ***D ≈ RW***, (1) a response profile matrix ***R*** ∈ R*m*×*k* that characterizes the response of each of *k ≪ n* components to all *m* stimuli, and (2) a component by unit weight matrix ***W*** ∈ R*k*×*n* that expresses the contribution of each unit to every component’s response. This approach is intrinsically axis-dependent. We performed the same factorization approach on high-dimensional responses in the human ventral visual pathway in the NSD. For every DCNN component, we then computed its correlation with the top five neural components, which we had previously shown to be selective to faces, scenes, bodies, text and food respectively [19].

For every neural component, we found a highly correlated (0.44 0.80) DCNN component. We visualized the response profiles of each of these matched DCNN components across all stimuli and correlated the component responses with the salience ratings for their corresponding category in each of these stimuli (Fig. 5). These salience ratings were collected in an independent behavioral experiment to quantify the noticeability of each of the five visual categories within individual stimuli [19].

**Fig. 5.**
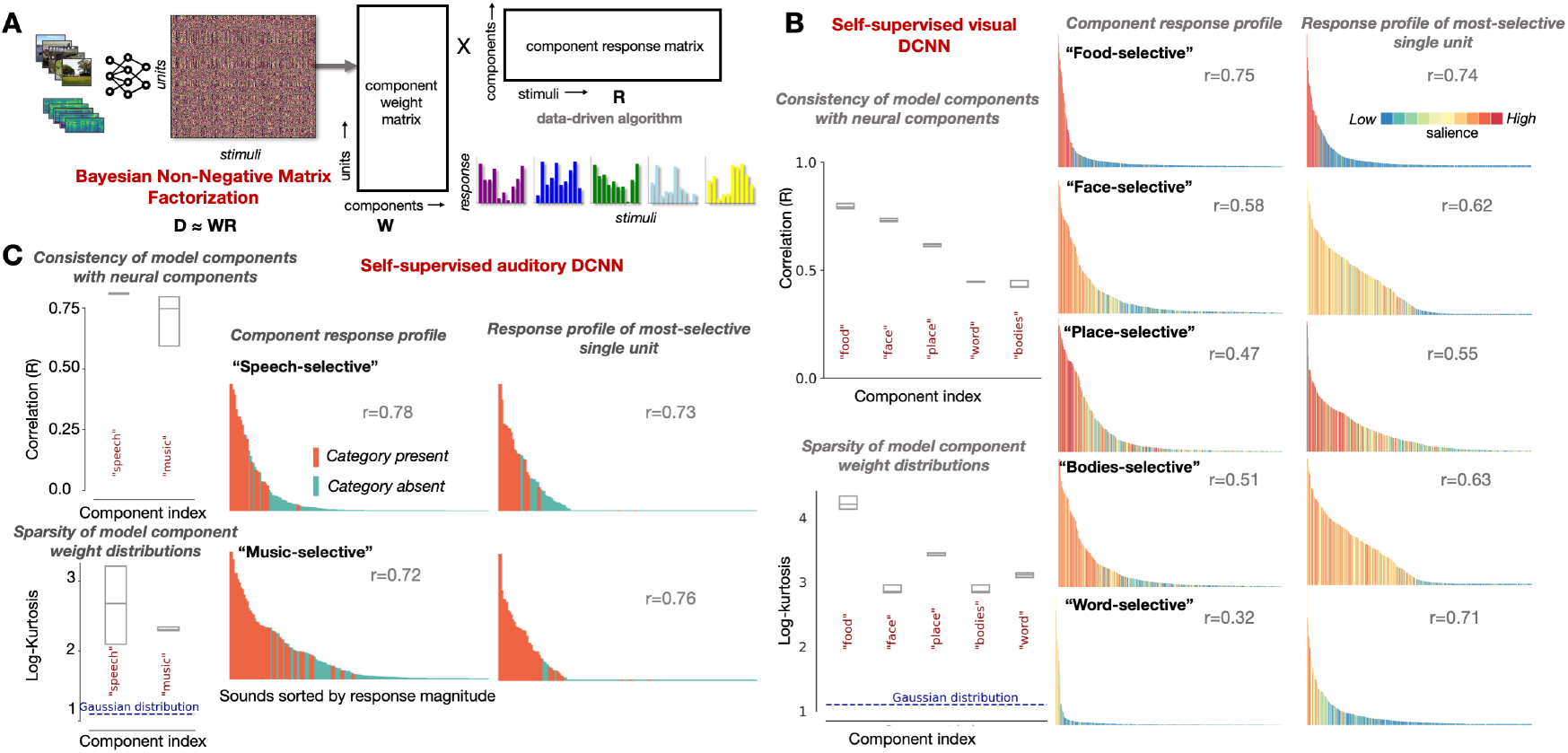
What are the preferred axes in brains and networks? (**A**) Overview of the factorization approach applied to derive the dominant components of DCNN representations. Bayesian non-negative matrix factorization was used to decompose DCNN activations as a product of two lower-dimensional matrices: (1) a response profile matrix characterizing the response of each component to all stimuli and (2) a component-by-unit weight matrix that expressed the contribution of each unit to every component. **B**) Left panel: (top) correlations between the top 5 visual DCNN components, sorted by their consistency with neural components, and their corresponding neural counterparts; (bottom) weight distribution sparsity (*κ*4) of these top 5 DCNN components across units. Middle panel: response profile of each component across the 515 images viewed by all NSD participants. Each bar corresponds to an image, with the color denoting the behavioral salience rating for the preferred category of the component (e.g. the salience of faces for Component 2). Correlation (Pearson’s R) between the magnitude of component responses and salience ratings for the preferred category of each component across all 515 images is reported in the insets. Right panel: response profile of the single unit that is most selective for the respective category. **C**) The same analysis as **B**, but for auditory DCNN components. Correlation (Pearson’s R) of component responses to all 165 sounds with a binary vector indicating whether the preferred category of the component (music/speech) was present/absent in the stimuli is reported in the insets. Boxplots reflect variability over multiple iterations of the NMF.

This analysis revealed that the DCNN component responses were each indeed highly selective to a different category. Are these category-selective component responses indicative of distributed responses among all units, or are they driven by a few specific units? To answer this, we next turned our attention to the component-by-unit weight matrix and computed the kurtosis of the unit weight distributions for each component. As shown in Fig. 5B (left, bottom), unit weight distributions were highly kurtotic, relative to a Gaussian, indicating a peakier, heavy-tailed distribution skewed toward higher values, suggesting that the components were indeed sparse and received contributions from only a few units, just like the neural components. We also evaluated whether there were single units in the DCNN selective for each of these five categories. This analysis revealed single units whose responses were highly correlated (*R* 0.44 0.80) with each of the categories, respectively (Fig. 5B, right). These findings are consistent with previous reports of category selectivity in DCNNs [49, 50], but extend previous findings by showing that category selectivity emerges in a hypothesis-neutral fashion as the dominant tuning in DCNNs, and in showing that these dominant responses are matched to those in the brain.

In the auditory self-supervised DCNN, the same factorization approach yielded an analogous result: sparse components highly selective for music and speech (Fig. 5C), again reminiscent of the category-selective tuning previously reported in the auditory cortices [37]. Analysis of single DCNN unit responses further reveals that some single units themselves exhibit selectivity for these complex high-level categories (Fig. 5C, right), consistent with one recent report of music selectivity in audio-trained DCNNs [51]. Control experiments showed that the observed selectivity for high-level abstract categories like music or speech was not explained by generic acoustic features, such as spectrotemporal modulations (see Supplementary Fig. S11).

Are these aforementioned categories more prominent in the tuning functions of the native axes than in arbitrary axes? To find out, we computed the number of units selective for each of these categories in both the native and arbitrary axes. As in previous analyses, we generated a sequence of rotation matrices that interpolate between a uniformly-drawn random rotation matrix and the identity matrix (no rotation). We determine the count of category-selective units in each of these representational bases for both sensory cortical representations and self-supervised DCNN representations. To compute selectivity, we contrast the response of each unit to stimuli containing the category in question against all other stimuli with a t-statistic (following the standard approach of functional localizers in neuroimaging experiments), comparing it to a fixed t-value threshold to identify the number of selective units. We used an image set encompassing text, bodies, faces, and places [52] and the 165 sounds from the auditory dataset [37].

As shown in Fig. 6, category-selective tuning is most pronounced in the native axes, with deviations from these axes consistently reducing single-unit selectivity. This result provides evidence that category-selective tuning is a hallmark of the privileged axes found in sensory systems and models. We note that the alignment of representational axes in penultimate DCNN layers with human-interpretable categories emerges independently of DCNN task performance, given that arbitrary rotations of the axes preserve downstream task performance.

**Fig. 6.**
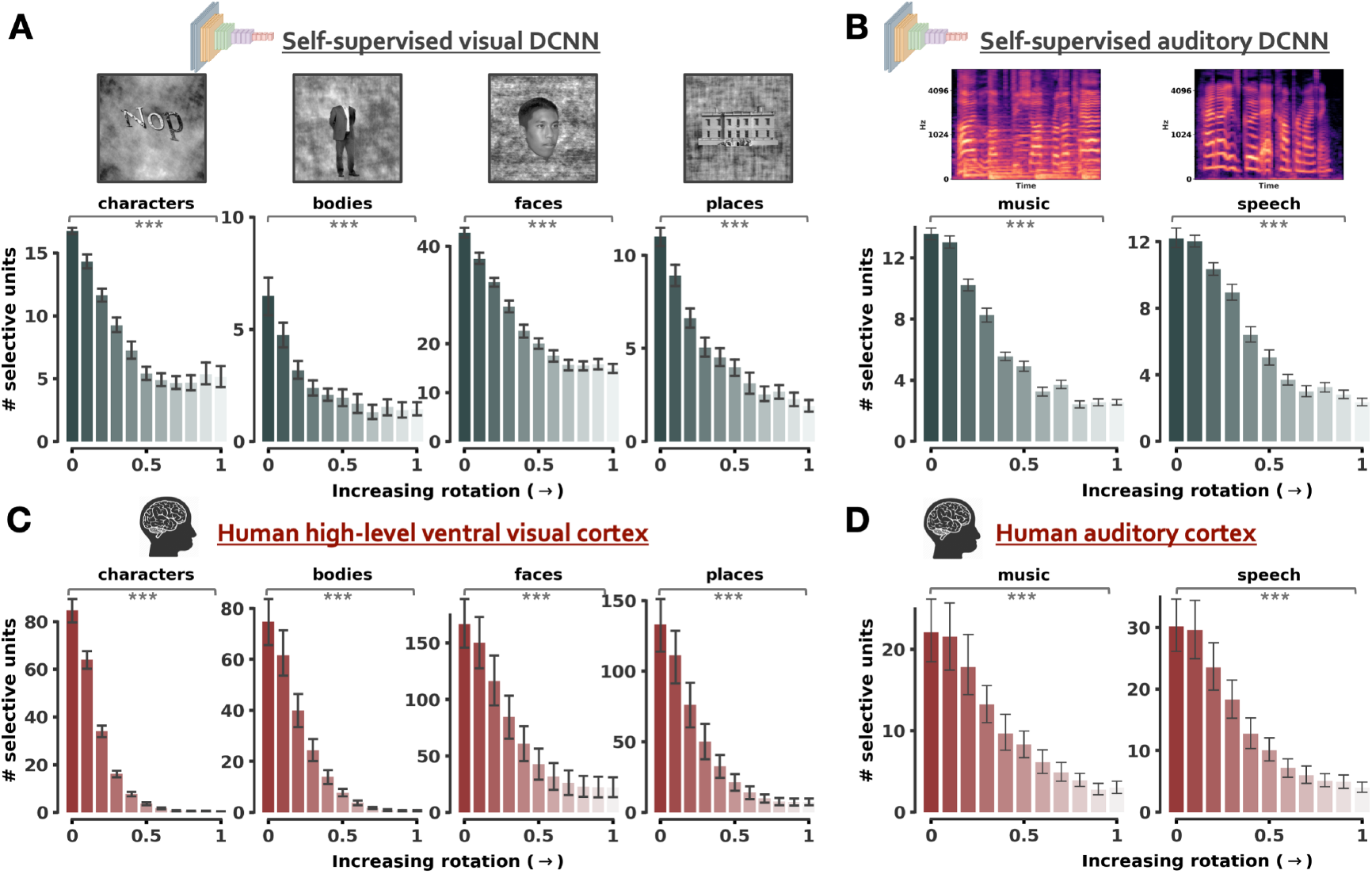
Category-selectivity is an axis-dependent phenomenon. (**A**) Number of single units in a self-supervised visual DCNN selective for each of four categories: characters, bodies, faces, places. Here, and in **B**, selectivity is assessed via t-statistics, contrasting responses to stimuli from the chosen category against others, mirroring standard functional localizer procedures. (**B**) Number of single units in a self-supervised auditory DCNN selective for music (left) and speech (right). Spectrograms of example stimuli for these categories are presented above. (**C**) and (**D**) show analogous plots for neural representations in the human high-level ventral visual cortex and auditory cortex, respectively. Here, the unit of analyses are single voxels.

The emergence of a sparse category-selective tuning structure in neural networks sheds light on their origins in the brain. While prevailing theories often attribute category selectivity in brains to downstream behavioral pressures and the ecological significance of these categories, our findings show that category selectivity can arise without any of these explicit pressures. Its presence in self-supervised models that are not optimized for a behavioral task indicates that category-selectivity can arise simply from the statistics of our natural input and optimization for contrastive objectives [49, 50], potentially in conjunction with particular architectural features like nonlinearities.

### Privileged axes dissociate neural architectures

To investigate whether privileged axes depend on the type of architecture underlying a representation, we extend the previous analyses to the transformer model architecture. The transformer has recently emerged as another promising model class that can successfully capture neural activity and human behavior across vision and audition [53–55]. Transformers do not contain convolutions but instead leverage parallel self-attention mechanisms to effectively capture contextual information by allowing every position in an input sequence (e.g. a patch in an image) to attend to every other position. Each transformer layer further contains a feed-forward neural network, also known as the Multi-layer Perceptron (MLP) block, that operates independently on each position and introduces non-linearity to the model through its activation functions (often, a softened nonnegativity constraint like GeLU). This complements the self-attention mechanism by enabling more complex transformations, equipping the transformer to model a rich set of functions.

Recent studies have revealed that transformer architectures achieve comparable performance to deep convolutional neural network architectures in predicting neural responses throughout the sensory cortices [56, 57]. Here, we ask whether transformers also exhibit axis alignment with the brain. We extracted visual representations from two vision transformers trained on ImageNet with category labels: (i) a standard ViT architecture (ViT) [54], and (ii) a hybrid model where the input sequence to the ViT is formed from intermediate feature maps of a regular ResNet50, henceforth denoted as ViT-RN50 [54]. We also extracted sound representations from two Auditory Spectrogram Transformers (AST) trained on the AudioSet corpus with category labels and initialized with different random seeds [55], henceforth denoted as AST-1 and AST-2. For each of these networks, we extracted the penultimate representation (output of the final attention block) as well as the representation in the last feedforward layer with a nonlinear GeLU activation function, labeled ‘GeLU’. We then compared each of these DCNN/transformer representations to the corresponding sensory areas using the most widely used brain-model similarity metric: linear predictivity after training a linear model that maps DNN activations to brain activity. We contrast this standard metric with our measures of axis-alignment.

Although DCNNs and transformers exhibited comparable linear predictivity (as expected given previous results), they differed markedly in the extent to which their axes were aligned to those of the brain (Figure 7A-C, right). The representational axes of DCNNs were much better aligned to those of the brain than were those of transformers. This trend held true both for macaque IT recordings and for human ventral visual stream fMRI measurements, and for human auditory cortical fMRI measurements. Further, in the visual domain, the DCNN-brain axis alignment was close to the estimated noise ceiling (as measured by inter-subject axis alignment). We next asked whether the representational axes of transformers exhibit a non-random relationship with sensory cortical representations depite being worse than that of DCNNs. To address this, we again computed the axis-dependent alignment as a function of axis rotation. While the basis of the penultimate layer was nearly indistinguishable from an arbitrary basis by its alignment with neural representations in both audition and vision, the GeLU-layer axes were significantly more aligned than an arbitrary basis. This result is consistent with the idea that nonlinearities are important for generating brain-like privileged axes (because the penultimate layer lacks an ensuing nonlinearity).

**Fig. 7.**
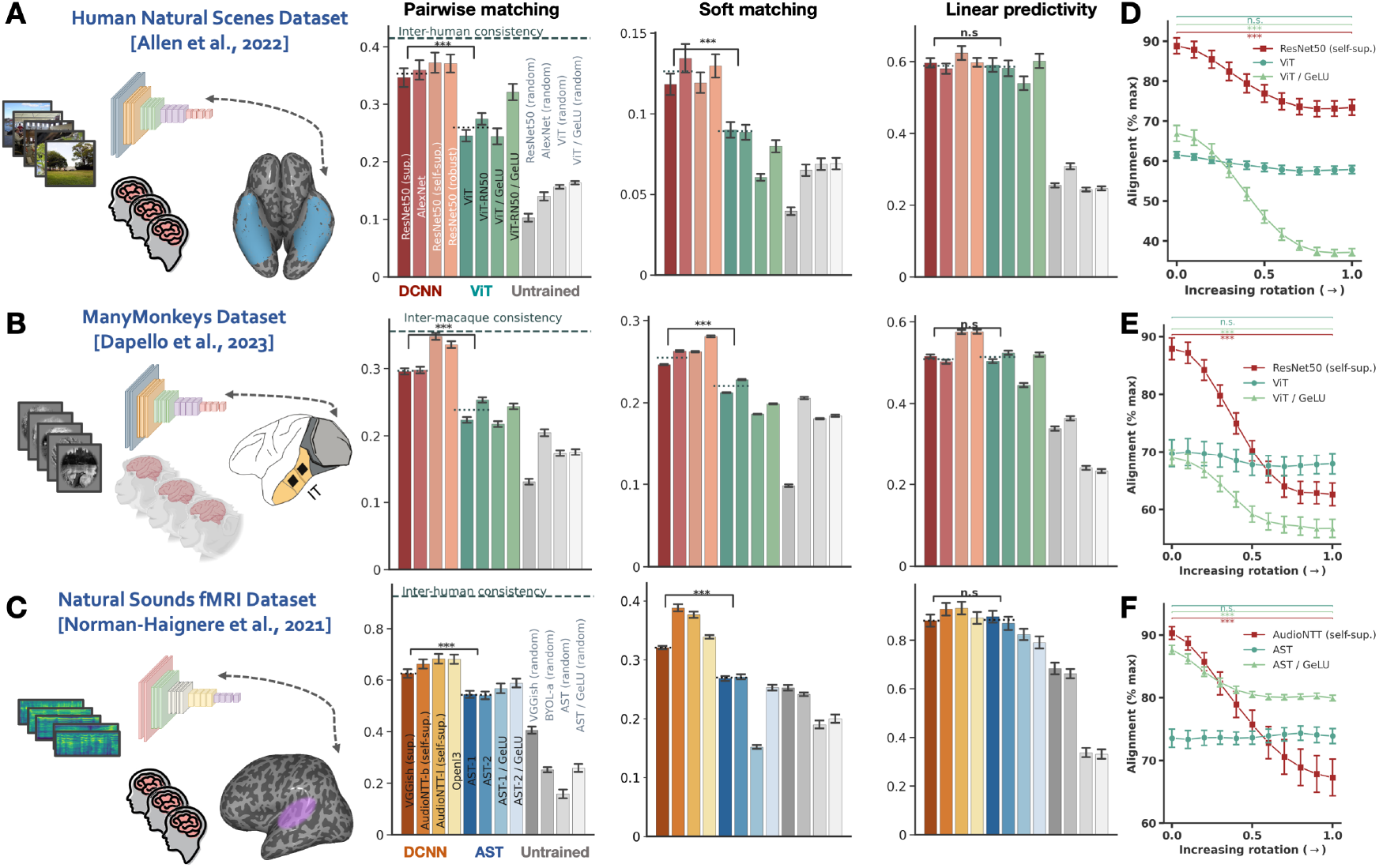
Axis-dependent alignment methods better discriminate fit of computational models to the brain. (**A-B**) Alignment of various visual deep convolutional neural networks with neural representations in the human ventral visual stream and macaque IT respectively, examined using two measures of axis-alignment (the pairwise matching correlation, left, and the soft matching correlation score, middle) and linear predictivity. Here and in **C**, left plots display inter-individual consistency for the pairwise matching correlation.(**C**) Alignment of auditory DCNNs with neural representations in the human auditory cortex, assessed using the same three metrics. (**D**) Pairwise matching correlation between various artificial neural network and brain representations as a function of model representation axis rotation (*α*), expressed as a proportion of the maximum possible value obtainable across all axis rotations, determined using the ‘Optimal Axis Alignment’ procedure described in the text. Here, the model representations include penultimate layers of self-supervised DCNNs (architectures ResNet50 and AudioNTT), transformers (ViT and AST) and the final GeLU layers of transformers (ViT / GeLU and AST / GeLU).

Altogether, these results show that by adopting a more stringent measure of similarity, our axis-alignment metric better discriminates which artificial network models best fit the brain.

### Computational implications of privileged axes

Given that axis rotations maintain the information content and population geometry of a representation, why do high-level sensory cortical and DCNN representations exhibit specific, non-random axes? We hypothesized that the representational form employed by the brain could affect efficiency, downstream wiring costs and the capacity for few-shot generalization under anatomical and physiological constraints.

#### Efficiency

We operationalized efficiency as economy in the number of neurons active in response to a stimulus. To test whether the native axes result in more efficient representations, we compared the lifetime sparseness of individual units in DCNN and cortical representations using the native basis, and compared it to that for uniformly-drawn random bases.

In both vision and audition, both brain and model native axes produced representations with higher lifetime sparseness than rotated axes, as measured with kurtosis (Fig. 8A) as well as an alternative measure of lifetime sparseness proposed by Vinje and Gallant [58] (Supplementary Fig. S12). This effect is also evident in the response distribution of units in the native axes, which had a more pronounced heavy-tailed appearance compared to units in an arbitrary basis (Fig. 8A left, top vs. bottom).

**Fig. 8.**
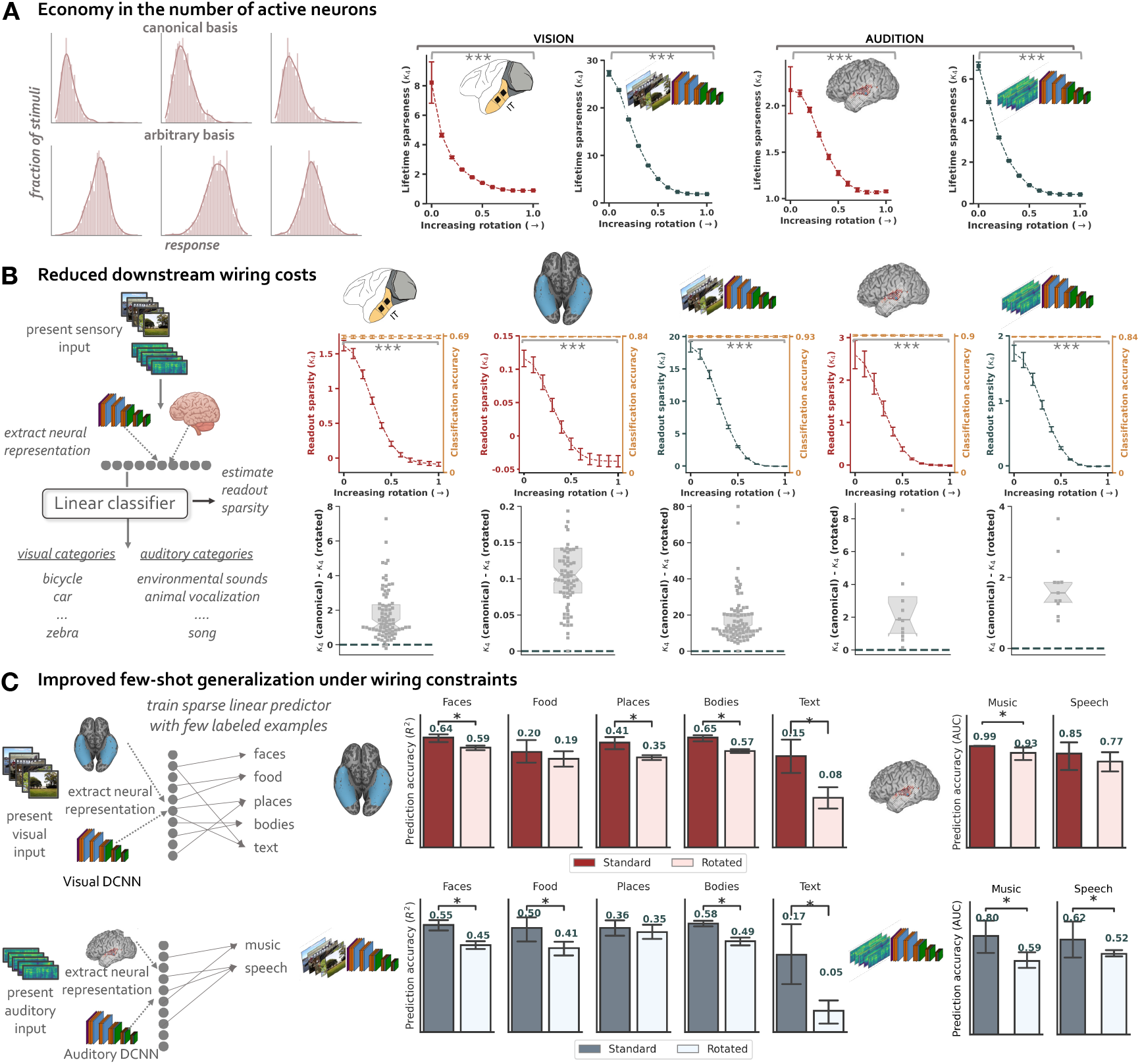
Computational implications of privileged axes. (**A**) Left panel: distribution of responses to natural stimuli for 3 neurons in the native basis (top) and 3 in the rotated basis (bottom) using the Macaque-NSD dataset. Right panel: lineplots show the mean lifetime sparseness, computed as kurtosis, across all units in the representation at different axis rotations. From left to right, the representations include the macaque IT neural responses (from the Macaque-NSD dataset), self-supervised visual DCNN activations, human auditory cortex responses (ECoG) and self-supervised auditory DCNN activations. (**B**) Left panel: schematic illustrating the procedure for estimating downstream readout sparsity. Linear decoders are trained on top of neural representations to predict stimulus categories. The sparsity of the learned readout weights are calculated as kurtosis. Right panels, top: Lineplots show the mean readout sparsity, computed as kurtosis, across all categories in the representation at different axis rotations. Right panels, bottom: Histograms depict the distribution of the difference in sparsity between the native and rotated axes across all categories. From left to right, the representations include the macaque IT neural responses, human ventral visual stream responses, self-supervised visual DCNN activations, human auditory cortex responses and self-supervised auditory DCNN activations. (**C**) Procedure for estimating the few-shot generalization performance of each representation under sparsity constraints. We train sparse linear readouts (using *l*1-regularized linear regression) on top of neural representations to predict the salience/presence of each category in the stimulus using only few labeled examples. Left panel shows the few-shot generalization performance of the human ventral visual cortex representation (top) and the self-supervised visual DCNN representation (bottom) for the regression task (i.e., predicting category salience), estimated using *R*2 for each of the 5 visual categories. Right panel shows the performance of the human auditory cortex representation (top) and the self-supervised auditory DCNN representation (bottom) for the classification task (i.e., predicting whether the category is present or not), estimated using *AUC − ROC* for each of the 2 auditory categories.

#### Downstream wiring costs

We next hypothesized that the axes in the brain are those that reduce wiring costs. Due to space constraints on the number of possible connections between neurons in two regions, a neuron in one region likely cannot connect to every neuron in another region, such that its readout from that region might need to be driven by a small subset of neurons. We asked whether the axes found in the brain are those that facilitate a sparse readout of behaviorally relevant information (e.g. object category labels) from patterns of neural activity.

We trained linear decoders on representations from DCNNs and sensory cortices, aiming to predict the category labels of different stimuli from their evoked representations. For visual representations, we used NSD images for extracting both model (here, the self-supervised visual DCNN) and brain representations (macaque IT and human VVC). The decoder classes were the 80 object categories in the MS-COCO dataset. Since an image in the NSD can have more than one object category, we set this up as a multi-label classification task and quantified the classification performance for each category label with the Area under the Receiver Operating Characteristic (AUC-ROC) measure. As shown in Fig. 8B, all model and sensory cortical representations perform well on this decoding task (AUC-ROC ~ 0.69-0.93). This decoding accuracy is not affected by the change of basis, since arbitrary rotations preserve the information that can be linearly accessed in a representation. We then assessed how sparsely the category information is ‘read out’ in each basis by estimating the kurtosis of the readout weight distribution of each category over all units. Results revealed that the native basis yields a much sparser readout than an arbitrary basis, and even small deviations from the native basis lead to notable reductions in readout sparsity. This reduction is not dominated by a few select categories, but almost every category is read out more sparsely in the native basis.

Analogous experiments in the auditory domain yielded similar results. Here, we trained linear classifiers on intracranial cortical responses and self-supervised DCNN activations to natural auditory sounds. The classifier decoded the category labels of the sounds, which spanned eleven human-interpretable categories (e.g. environmental sounds, human vocalizations, song etc.). As in the visual domain, a change of basis preserved the decoding accuracies but dramatically decreased the sparsity of the classifier weights. These results indicate that object category decoding can be implemented with lower wiring cost using the axes found in the brain. Whether the same conclusions hold for the within-category information presumably contained within many of these neural populations is an important question for future research.

#### Capacity for few-shot generalization under wiring constraints

We also hypothesized that privileged neural axes might have consequences for generalization, particularly impacting sample efficiency during learning of downstream tasks under biological constraints that might limit the number of neurons accessible for read-out. Specifically, we posited that the axes privileged in the brain could enhance generalization capabilities for the categories whose selective responses are manifested in those privileged axes in downstream linear readouts during constrained learning scenarios, such as few-shot learning and enforced sparsity on the readout. To test this, we trained sparse linear decoders on top of neural representations in both native and arbitrary bases. A sparsity constraint, meant to mimic anatomical constraints on the number of inter-region connections, was enforced on the decoder through an *l*1 penalty. The *l*1 penalty also makes the generalization performance of the decoder axis-dependent. A few-shot learning scenario was instantiated by training the linear decoders using only a a very small fraction (10%) of labeled stimulus-category pairs. In the visual domain, separate sparse linear regressors were trained to predict the presence of each of the five categories in an image (quantified using salience ratings): faces, places, bodies, text and food. For auditory representations, two distinct sparse linear classifiers were utilized to discern the presence of music or speech in a sound waveform. For each candidate representation and basis (native vs. arbitrary), we then computed the generalization performance of the trained sparse linear decoders on held-out stimuli. Across most categories, the generalization performance of a representation in its native basis significantly exceeded its performance in an arbitrary basis. This pattern was consistent in both neural network and sensory cortical representations, and was observed across both visual and auditory domains.

Why do privileged axes facilitate few-shot learning? Intuitively, generalization error in a decoding model comprises a bias component, stemming from the decoding model’s simplifying assumptions, and a variance component, arising from fluctuations in the decoder weights based on the training set. Imposing a sparsity constraint will reduce the variance component across all axes by narrowing the hypothesis space. However, the bias component is then significantly influenced by specific axes. For many axes, the sparsity penalty might exclude the actual solution from the hypothesis space, increasing bias and, in turn, the generalization error. See Methods for a more complete description of this theoretical proposition.

### Privileged axes exist beyond the sensory cortices in higher-order neural systems

Are privileged axes exclusive to sensory domains, or could neural systems supporting higher-level cognition also privilege certain coordinate systems over others? Here, we consider one such high-level brain region: the human language cortex–associated with language comprehension and production.

To extract language cortex representations, we analyzed fMRI activations measured in nine participants as they read 384 sentences, each representing a unique stimulus condition, from short passages covering a wide variety of topics [59]. We calculated representational alignment for each pair of human subjects within this dataset, both in the native axes and across a sequence of arbitrary axes derived from axis rotations. Even in the language areas, alignment was significantly higher for the native axes than any arbitrary axes (Fig. 9). These results raise the possibility that the phenomenon of privileged axes may be ubiquitous throughout the cortex. We leave a through characterization of these privileged axes as well as the parallel analysis of computational models of these high-level cognitive abilities for future work.

**Fig. 9.**
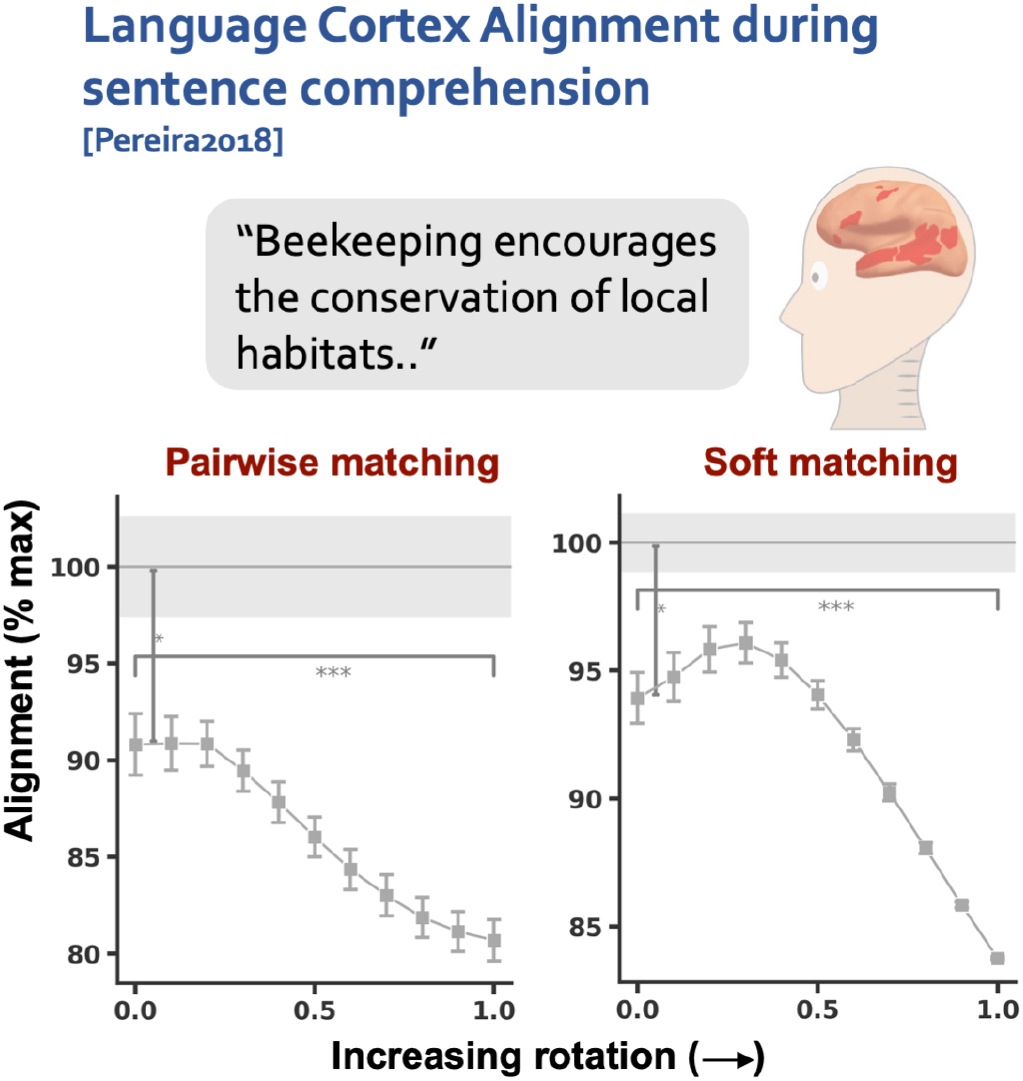
Alignment between pairs of conspecifics neural responses in the human language cortex during sentence comprehension, measured using the pairwise matching (left) and soft matching correlation (right) scores and examined as a function of axis rotation (*α*) in one participant’s representation. Alignment is expressed as a proportion of the maximum possible value obtainable across all axis rotations, determined using the ‘Optimal Axis Alignment’ procedure described in the text. Error bars are SEM over multiple splits of the data. Significance tests compare the alignment values at no rotation (*α* = 0) with those at full rotation (*α* = 1), as well as with the optimal alignment values observed across all rotation angles.

## Discussion

How is information coded in the brain? To answer this question, neuroscientists have increasingly adopted methods that analyze the geometry of the population code that are blind to the actual tuning of neurons. But are neural tunings in fact arbitrary and irrelevant, or might they matter, privileging some representational axes over others? We developed methods to probe for privileged representational axes in biological and artificial neural networks, and applied them to multiple types of neural data from diverse brain systems, and to DCNNs trained on natural sensory stimuli. We found that representational axes were consistent between individuals, between artificial neural networks varying in architecture and learning objectives, and between brain systems and artificial neural networks trained on the same modality. These results indicate that representational axes in neural systems are not arbitrary, and can arise in artificial systems with none of the priors and meanings assigned to these axes by humans. We further found that the privileged axes used in the brain and DCNNs are those that yield sparse representations and that aid read-out and generalization by downstream classifiers. Finally, our metric of axis alignment also distinguishes the fit of models to the brain that are not well discriminated by standard metrics.

### Privileged axes in brains

A key contribution of the present work is to develop and apply quantitative measures to formally test whether the representational axes in different animals/subjects are more similar than would be expected from arbitrary bases, and on the other hand whether they approach the maximum alignment possible. Our findings are to our knowledge the first quantitative evidence that high-level sensory cortices possess preferred axes for encoding natural stimuli. Our findings do not dismiss the utility of population-level analyses in understanding the neural code, but rather propose that such analyses should not ignore neural tuning functions. We found that among the large set of population codes with identical geometry and content, only some are empirically observed. We later discuss potential mechanistic reasons for why certain assignments of representations to individual neurons are favored, along with their computational implications.

### Privileged axes in models

Previous research has offered hints of privileged axes in artificial neural networks, such as the emergence of edge detector units or units selective for specific semantic concepts across a variety of networks [40, 60–66]. Such observations suggest a possible alignment of unit tuning curves beyond just the alignment of representational content. However, these qualitative observations of tuning similarity leave it unclear whether the alignment of tuning functions across networks exceeds what would be expected by chance. Our approach allows this comparison, and reveals that the axis alignment across different models far exceeds that observed in arbitrary bases, indeed approaching the maximal possible alignment achievable through any axis rotation. While the existence of privileged axes in artificial neural networks doesn’t guarantee that these axes will be interpretable to humans, our experiments suggest that that in many cases they are. At the very least, our findings provide a clear rationale for focusing on the native axes, rendering interpretability analyses of these bases not just convenient, but well-founded.

### Aligned privileged axes in brains and models

A large body of work supports the idea that different computational architectures often end up learning similar representations (known as the universality, or ‘convergent learning’ conjecture in machine learning) [66–68]. Over the last decade, it has been suggested that this universality also extends to comparisons between artificial and biological networks, with task-optimized neural networks argued to reproduce aspects of biological representations in multiple domains, including vision [27, 69], audition [57, 70] and language [71–75]. This representational convergence across brains and models is taken to reflect the strength of behavioral constraints and the statistical regularities of our natural world in narrowing down the solution set for a nervous system. However, the results supporting this convergence primarily rely on measures of representational content that are invariant to axis rotations, most commonly neural predictivity under a linear mapping model and representational similarity analysis. Our results show that the convergence of biological and artificial neural systems goes beyond representational content to encompass representational form. This conclusion is consistent with several recent papers that also note alignment between the representational units of the visual cortex and DCNNs [49, 76, 77]. The comparison against arbitrary bases in our work underscores the nontrivial nature of this similarity.

What causes brains and DCNNs to converge onto similar axes? Learning objectives by themselves should not result in preferred axes (because axis rotation preserves task performance for linear readouts). Rather, they must interact with the neural architecture to yield consistent axes. In artificial neural networks, particularly those employing ReLU nonlinearity, we can understand the representational invariances and the mechanisms leading to the emergence of privileged axes through theoretical constructs like the set of operations that commute with the nonlinearity (*GReLU*). We speculate that the privileged axes arising in the brain might be similarly causally linked to the nonlinear activation functions of neurons, and that such non-linearities could explain the axis alignment between biological and artificial neural networks.

### Interpreting privileged axes

Given that brains and ANNs share privileged representational axes, what are they? To address this question in a data-driven rather than ad hoc fashion, we used a sparse factorization algorithm to pick out the dominant tuning functions in both deep neural network representations and the corresponding neural systems. In both cases the predominant tuning functions were category selective for the categories previously identified in prior studies of neural tuning (faces, places, bodies, and words for visual networks and brain regions, and speech and music for auditory networks and brain regions), despite not being constrained in any way to be interpretable or to conform to presupposed hypotheses. A recent preprint also described category-selective tuning in networks trained on visual self-supervised learning objectives [49, 76].

Our study further shows that brain-like category selectivity in individual network units (i) is not confined to visual processing, but also extends to auditory models, (ii) is a dominant feature of high-level DCNN representations, as the specialized categories emerge spontaneously from a data-driven decomposition of network activations, and (iii) is not an inevitable result of any high-level representational space, since rotating the axes of the representation preserves the content but eliminates selective responses. Our findings show that category selectivity can emerge merely from the statistics of the natural world in conjunction with neural architecture constraints, without specific priors for any of these categories (see also [49]). This is in contrast to the conventional view of selective responses, which attribute the development of selective responses in the cortex directly to computational advantages afforded by selective responses, e.g. augmenting storage capacity in associative memories, overcoming the superposition catastrophe, facilitating post-perceptual processing etc. [78–80].

### Computational implications of privileged axes

Representations are best understood in the context of the computations that transform them for use in downstream tasks [81]. While there have been many studies comparing the downstream impact of different kinds of representations, for instance analyzing the effect of the dimensionality of a representation or how representations with mixed selectivity compare to representations with ‘pure’ selectivity [82], little attention has been paid to the impact of representational axes. We proposed several respects in which representational axes could impact computation, all related in some way to sparsity. First, axes affect the efficiency of the neural code. We found that axes in both biological and DCNNs are characterized by heavy-tailed (kurtotic) response distributions when processing natural stimuli. High kurtosis is one measure of lifetime sparseness, and is associated with efficient (non-redundant) representations [83]. This finding is consistent with previous reports of high lifetime sparseness in early and high-level visual areas [58, 78, 84, 85]. Second, the axes affect downstream wiring costs. Readouts in the brain are likely to have to be restricted to small subsets of all possible neurons in order to avoid prohibitive volumes of brain tissue. We found that the axes of both biological networks and DCNNs facilitate sparser readouts of category information compared to arbitrary bases. This finding may relate to a recent study of neural responses in primate prefrontal cortex during a working memory task, revealing how neural implementations of representational geometries influence wiring costs [86]. Third, axes affect the capacity for generalization. If connections are constrained to be sparse, the generalization of any readout of a representation is dependent not only the representation’s geometry but also on the tuning of representational units. We found that the axes observed in the brain support better few-shot generalization under sparse linear readouts compared to arbitrary bases. Our theoretical analysis suggests that any task that relies on contributions from a small subset of neurons will show better few-shot generalization if the representational axes are those in which natural stimuli are sparsely represented. A similar relationship between disentanglement and improved generalization in the context of multi-task learning was also noted in recent theoretical work [87].

### The importance of axis-dependent alignment measures in comparative analyses

Given our finding that systematic neural tuning is a basic fact of brains, it requires explanation, and metrics that are blind to this phenomenon are incomplete at best. Our test of axis alignment fills this gap, and enables better discrimination between models in terms of their fit to neural data. For instance, we found that DCNNs exhibit greater similarity to biological sensory systems than transformers when evaluated via axis alignment, but not when using axis-independent measures. Further, our finding of one-to-one mappings between units in networks and units in brains constitutes evidence of a more literal and stringent kind of similarity between networks and brains (see also [50]), more strongly supporting the use of these networks as models of the brain. These results indicate that measures of axis alignment, along with other recent model-brain comparison metrics, should be added to commonly employed batteries of model-brain comparisons.

### Limitations and future directions

Our work is subject to several limitations. First, assessing the prevalence of privileged axes in the brain is limited by existing neural activity data, which are currently limited in either coverage (electrophysiology) or spatial and temporal resolution (fMRI). Thus it is possible that other representational axes may exist that are less consistent across participants or between brains and ANNs. Second, our reliance on gradient-based optimization procedures to derive the optimal alignment could present limitations in certain situations and for certain models (by working less well due to local optima). Third, while our work suggests that nonlinearities could play a role in determining tuning functions, it remains unknown which nonlinearities produce brain-like privileged axes in computational models of neurobiological systems. Recent work has provided theoretical and empirical evidence that in the presence of biological constraints on neurons like non-negativity (as routinely employed in modern DCNNs via ReLU operations) and sparsity, single neurons become aligned to axes of data generative factors [88]. However, whether this happens for realistic natural sensory input is an open question. Studies of how experience with varying sensory diets shapes the tuning functions in in-silico models could provide insight. Finally, the concept of axis alignment in the brain is scale-dependent. We focused here on a meso-scale level of analysis, considering representations within modality-specific cortices and networks and finding privileged axes in all cases we examined. But this work leaves open the question of whether there is a scale of analysis at which representations transition from privileged axes to arbitrary axes. Our preliminary analyses of axis alignment within category-selective regions of the high-level visual cortex suggests that at this finer scale, voxel responses in conspecifics are no more aligned in the native basis than an arbitrary basis, providing hints that at this scale representations may not have privileged axes (see Supplementary Fig. S13). However, these conclusions are limited by the coarse resolution of (fMRI). To validate these initial observations, further research employing higher spatial resolution within category-selective patches is essential.

Perhaps most fundamentally, our work does not answer why biological and artificial networks have representational dimensions selective to some categories of input (e.g. faces, places, speech) but apparently not others. One speculative hypothesis is that the data generating causes in the natural world consist of sparsely interacting parts, and DNNs that perform (generalize) well learn to represent these parts in individual neurons in the presence of certain nonlinearities. This idea resembles the “Platonic representation hypothesis” [89] but takes it one step further to argue that not only do networks (and brains) tend to converge to extract on the causal structure of the world, but many of the key dimensions of the world will be reflected in neurons selective for that dimension.

## Methods

### Brain Activity Datasets

#### Natural Scenes Dataset

A detailed description of the Natural Scenes Dataset (NSD; http://naturalscenesdataset.org) is provided elsewhere [35]. Here, we briefly summarize the data acquisition and preprocessing steps. The NSD dataset contains measurements of fMRI responses from 8 participants who each viewed 9,000–10,000 distinct color natural scenes (22,000–30,000 trials) over the course of 30–40 scan sessions. Scanning was conducted at 7T using whole-brain gradient-echo EPI at 1.8-mm resolution and 1.6-s repetition time. Images were taken from the Microsoft Common Objects in Context (COCO) database [90], square cropped, and presented at a size of 8.4° x 8.4°. A special set of 1,000 images were shared across subjects; the remaining images were mutually exclusive across subjects. Images were presented for 3 s with 1-s gaps in between images. Subjects fixated centrally and performed a recognition task in which they were instructed to indicate whether they have seen each presented image at any point in the past. Informed consent was obtained from the participants and the study was approved by the University of Minnesota Institutional Review Board.

The fMRI data were pre-processed by performing one temporal interpolation (to correct for slice time differences) and one spatial interpolation (to correct for head motion). A general linear model was then used to estimate single-trial beta weights, with the HRF estimated for each voxel and the GLMdenoise technique used for denoising [91, 92]. In this work, we used the 1.8mm volume ‘nativesurface’ preparation of the NSD data and version 3 of the NSD single-trial betas (‘beta fithrf GLMdenoise RR’). Every stimulus considered in this study had 3 repetitions, and we only analyzed data in the four NSD subjects who had full 3 repetitions for each of their 10,000 images. To derive voxel responses to each stimulus we averaged single-trial betas after z-scoring every voxel within each scan session.

We selected ventral visual stream voxels by using the “streams” atlas provided in the native space of each subject with NSD [35]. Briefly, this ROI collection reflects large-scale divisions of the visual cortex into primary visual cortex and intermediate and high-level ventral, lateral and dorsal visual areas. These were manually drawn for each subject by NSD curators and were based on voxel-level reliability metrics. For this study, we extracted the ROI mask corresponding to the ‘higher-level ventral stream’ label. This ROI was drawn to follow the anterior lingual sulcus (ALS), including the anterior lingual gyrus (ALG) on its inferior border and to follow the inferior lip of the inferior temporal sulcus (ITS) on its superior border. The anterior border was drawn based on the midpoint of the occipital temporal sulcus (OTS). This parcel is very broad, and results in 7, 000 − 9, 000 voxels per subject.

#### ManyMonkeys Dataset

The ManyMonkeys Dataset, as described in [93] and [36], comprises neural recordings from the inferotemporal (IT) cortex of six rhesus monkeys, captured during a passive viewing task. This dataset features a collection of neural responses to a set of 640 high-definition images. Each image represents a 2D rendering of a 3D object model superimposed on a randomly generated background, with variations in the object’s position, size, and orientation. The diverse stimulus array includes images of bears, elephants, human faces, apples, cars, dogs, chairs, airplanes, birds, and zebras.

Neural activities were monitored through the implantation of two to three micro-electrode arrays in the IT cortex of the monkeys, accessing approximately 58 to 280 neural sites in each monkey. These arrays were strategically placed to sample a breadth of neural sites across the posterior-anterior axis of the IT cortex. Spike events were identified in the voltage traces based on a threshold-crossing event, specifically when the falling edge of the voltage signal dropped below three standard deviations of the baseline voltage fluctuations.

For each image presented, the response of a neural site was quantified as the average firing rate within a 70-170ms window following the image’s appearance.

#### Monkey-NSD Dataset

The Monkey-NSD Dataset, collected by Yoon Bai in Jim DiCarlo’s lab at MIT, comprises neural recordings from the inferotemporal (IT) cortex of one monkey to the shared 1,000 images viewed by all participants of the NSD. It encompasses a total of 116 neural recording sites, with 49 located in the posterior IT region and 67 in the anterior IT. For this particular dataset, an advanced spike-sorting procedure was employed to identify spike events, offering an enhancement in the reliability of the recorded data compared to traditional spike-thresholding methods.

#### Auditory Datasets

**fMRI** The fMRI data analyzed here was originally collected and analyzed by Norman-Haignere et al., (2015). We summarize the data collection procedure with slight modifications from the original study for brevity.

Ten individuals participated in the study, each undergoing two to three fMRI sessions, each lasting 1.5 hours. They were scanned while listening to a set of 165 natural sounds, each lasting 2 seconds, designed to broadly sample common auditory experience. The experimental design utilized a block structure with each 17-second block comprising five presentations of one sound clip separated by scanner acquisitions. Within each block, one sound was presented at a 7 dB lower intensity to enable a sound intensity discrimination task. Participants were instructed to respond to this quieter sound by pressing a button. Following each sound, a corresponding fMRI volume was acquired using sparse scanning techniques that avoid temporal overlap between auditory stimulus presentation and scanner acquisition sounds. This resulted in a total of eleven runs per participant, each including fifteen sound blocks interspersed with four silence blocks to facilitate baseline response measurement. The functional scans were performed on a 3T scanner equipped with a 32-channel head coil, with a repetition time of 3.4 seconds and an in-plane resolution of 2.1 x 2.1 mm, leading to a voxel size of 2.1 x 2.1 x 4 mm.

The acquired fMRI volumes underwent several preprocessing steps using FSL and custom MATLAB scripts. Motion and slice timing corrections were applied, non-brain tissue was removed, and the voxel time series were linearly detrended. Each functional run was co-registered to the participant’s anatomical volume. Subsequently, these volumes were mapped to vertices on the cortical surface reconstructed via FreeSurfer, and spatial smoothing was applied using a 2D Gaussian kernel with a full width at half maximum (FWHM) of 3 mm.

The average response of a voxel to each sound was computed by analyzing the acquisitions from the second to the fifth post-stimulus onset, discounting the first to account for hemodynamic delay. These averaged responses were expressed as percent signal change (PSC) relative to the response during silence blocks. The PSC data were then interpolated onto a 2 mm isotropic grid on the cortical sheet.

Adopting the criteria from Norman-Haignere et al. [37], we selected voxels within a defined anatomical region that included the superior temporal and posterior parietal cortex, whose response reliability for the auditory stimuli exceeded a criterion. This resulted in a dataset of 11,065 voxels across all participants, with an average of 1,106.5 voxels per participant (range: 799-1498).

**Electrocorticography** The human intracranial electrocorticography (ECoG) recordings were initially acquired for a separate study [48]. The stimulus set for this dataset comprised the same set of 165 natural sounds used in the fMRI study described above [37]. The study involved fifteen individuals diagnosed with epilepsy and implanted with subdural electrode grids for clinical purposes. Superior temporal gyrus (STG) exposure was notably greater in one hemisphere for each participant (eight with right hemisphere placement and seven with left hemisphere placement). The predominant electrode configuration featured a 2.3 mm diameter electrode with an interelectrode distance of 6 mm on temporal lobe arrays (with a 10 mm spacing on arrays positioned in the frontal, parietal, and occipital lobes, though these generally did not yield consistent sound-evoked responses). Two patients received implants with a denser grid arrangement (1 mm exposed diameter, 3 mm inter-electrode spacing). Additionally, one individual was fitted with stereotactic electrodes in lieu of ECoG grids. Participants completed a variable number of experimental runs, ranging from three to seven.

The selection of electrodes for the final analysis was guided by the reliability of the broadband gamma activity (70–140 Hz) elicited by the acoustic stimuli, specifically the split-half correlation between responses to odd and even presentations, which had to exceed 0.2. This criterion resulted in inclusion of 272 electrodes across all participants. The signals from these electrodes were subjected to a common-average referencing process, calibrated against the mean across all electrodes within each subject. To ensure the purity of this referencing, electrodes with abnormal 60 Hz power levels were omitted. Subsequent to this referencing, a notch filter was applied to attenuate 60 Hz electrical noise and its harmonics. Broadband gamma power was quantified by the signal’s envelope within the 70–140 Hz filtered range. The response of each electrode during a three-second window locked to sound onset was measured, with each auditory stimulus lasting two seconds. These responses were normalized to percent signal change from baseline (the response during silence) by calculating the deviation from, and then dividing by, the average activity in the 300 ms pre-stimulus interval. For comparative analysis with neural network models, we averaged the responses within the three-second post-onset window for each electrode to each stimulus, treating each electrode as a representational unit. We also break down the analysis of model-to-brain alignment in different temporal windows for this ECoG dataset and find that the raw alignment scores as well the difference in alignment of the auditory DCNN representation with auditory cortex representations at no rotation and full rotation of the DCNN axis peaks around 300 ms following stimulus onset (see Supplementary Fig. S10).

### Model selection

We compiled a set of models spanning different architectures and learning objectives that we could compare to brain data. We analyzed late model stages within each candidate model on the grounds that they provide a hypothesis of a neural implementation of high-level auditory or visual processing. Specifically, we analyzed the last convolutional layer for CNN models, and the last attention layer for transformer models. For CNN models, we averaged the activations across the spatial dimensions to derive a single response value per unit (channel/filter). For transformer models, we averaged across tokens.

In the visual domain, we investigated a set of *n* = 6 candidate models which included both deep convolutional and vision transformer architectures, all trained on ImageNet [38]. Table 1 provides an overview of the candidate visual models.

**Table 1.**
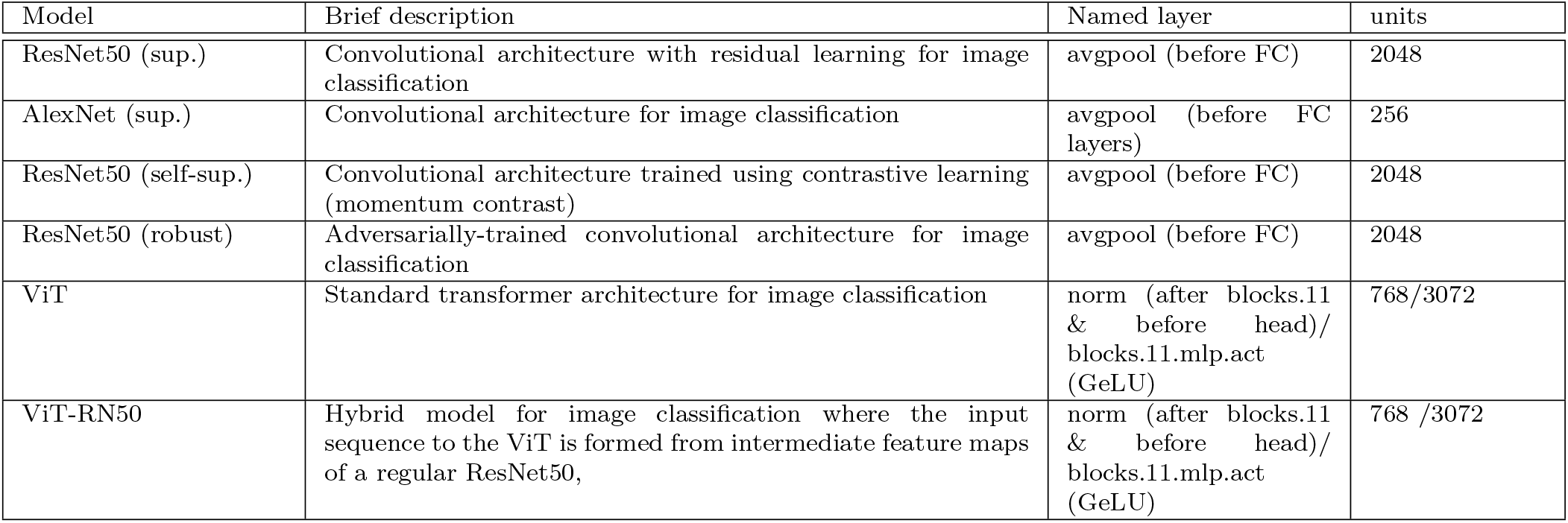
Overview of vision models.

| Model | Brief description | Named layer | units |
| --- | --- | --- | --- |
| ResNet50 (sup.) | Convolutional architecture with residual learning for image classification | avgpool (before FC) | 2048 |
| AlexNet (sup.) | Convolutional architecture for image classification | avgpool (before FC layers) | 256 |
| ResNet50 (self-sup.) | Convolutional architecture trained using contrastive learning (momentum contrast) | avgpool (before FC) | 2048 |
| ResNet50 (robust) | Adversarially-trained convolutional architecture for image classification | avgpool (before FC) | 2048 |
| ViT | Standard transformer architecture for image classification | norm (after blocks.11 & before head)/ blocks.11.mlp.act (GeLU) | 768/3072 |
| ViT-RN50 | Hybrid model for image classification where the input sequence to the ViT is formed from intermediate feature maps of a regular ResNet50, | norm (after blocks.11 & before head)/ blocks.11.mlp.act (GeLU) | 768 /3072 |

In the auditory domain, we again investigated a set of *n* = 6 candidate models which included both deep convolutional and transformer architectures, all trained on AudioSet [43]. Table 2 provides an overview of the candidate auditory models.

**Table 2.**
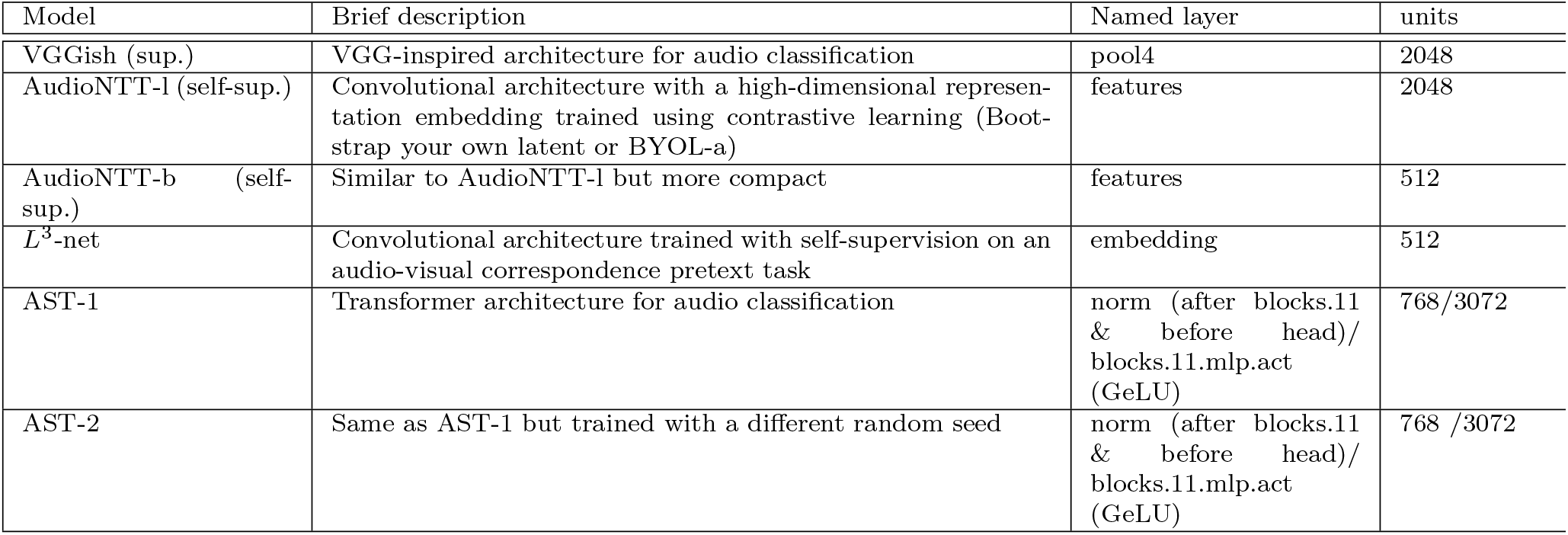
Overview of auditory models.

### Methodological framework for probing axis alignment

We propose a formal framework for assessing *whether* and *to what extent* a representational system (i.e., a brain region or DCNN) has a privileged representational axis.

Our framework entails three main steps. First, we propose two complementary measures of axis alignment. We use each to measure the alignment between two brains or two artificial neural networks or between a brain and a network. Second, we measure how this axis alignment changes when one of the two instances undergoes axis perturbations. Third, we compare the axis alignment under native and perturbed axes to the maximum alignment achievable under any axis rotations to assess the extent to which tuning functions expressed in the native axes are privileged.

All analyses of alignment were performed using training, validation, and test splits of data. When measuring alignment between two brains, two models, or a model and a brain, we used the training split to obtain the correspondence between representations, and the test split to measure the alignment between representations with independent data. When estimating the maximum possible alignment, we performed gradient descent using a training set, then used a validation set to determine the training epoch that gave the best alignment, and then used the test split to measure the alignment value.

#### Measures of axis alignment

Existing approaches for comparing representations focus on population-level similarity measures based on linear predictivity (as assessed using l2-regularized linear regression models) or RSA. These measures are insensitive to the axes (for instance, RSA is invariant to axis rotations) and hence cannot be employed to study axis alignment. To overcome this limitation, we propose two new measures for comparing the similarity between tuning functions in different representations.

##### Pairwise Matching score

Most analyses in the main text use a direct axis-dependent measure of pairwise similarity between units in different representational systems (the only exceptions are Figures 7 and 9, which also present the results for a Soft Matching score described below). Here, a unit in a biological representation corresponds to the smallest unit of brain measurement (e.g., a voxel in fMRI data, a single neuron in neurophysiology data), whereas a unit in an artificial neural network corresponds to a single “filter” in the convolutional layer of a DCNN or an activation unit in the MLP layer of a transformer model.

Consider matrices 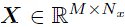 and 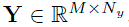, which represent the activations of *Nx* and *Ny* units respectively, for M instances. The Pairwise Matching score employs a bipartite semi-matching assignment algorithm [32] to find the alignment between these two representations, and proceeds in two stages:

- In the first stage, we find the optimal assignment matrix 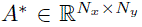 that maps each unit in ***X*** to a single unit in ***Y*** by optimizing a correlation-based objective. The optimization problem is defined as follows,

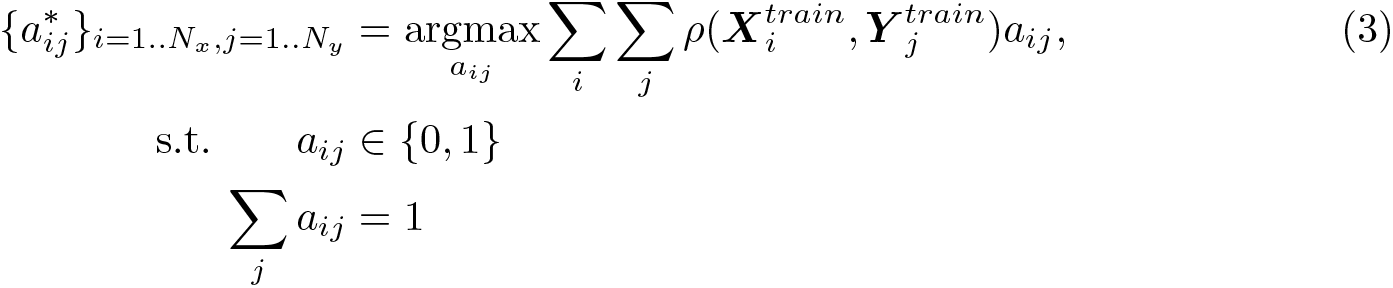 Intuitively, the optimization process aims to find the unit in ***Y*** that exhibits the maximum response correlation with each unit in ***X*** based on a training stimulus set. The constraint ensures that each unit in ***X*** is matched to exactly one unit in ***Y*** as shown in Figure 10A. As a result, some units in *Y* may remain unmatched after the assignment.
- In the second stage, using the optimal assignment matrix *A*∗ obtained from the first stage, we compute the correlation-based objective on held-out data. This entails calculating the correlation between each unit and its corresponding matched unit on the held-out data and then averaging these values across all matched pairs to derive the final alignment metric:

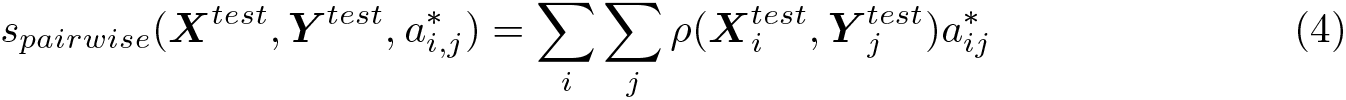

**Fig. 10.**
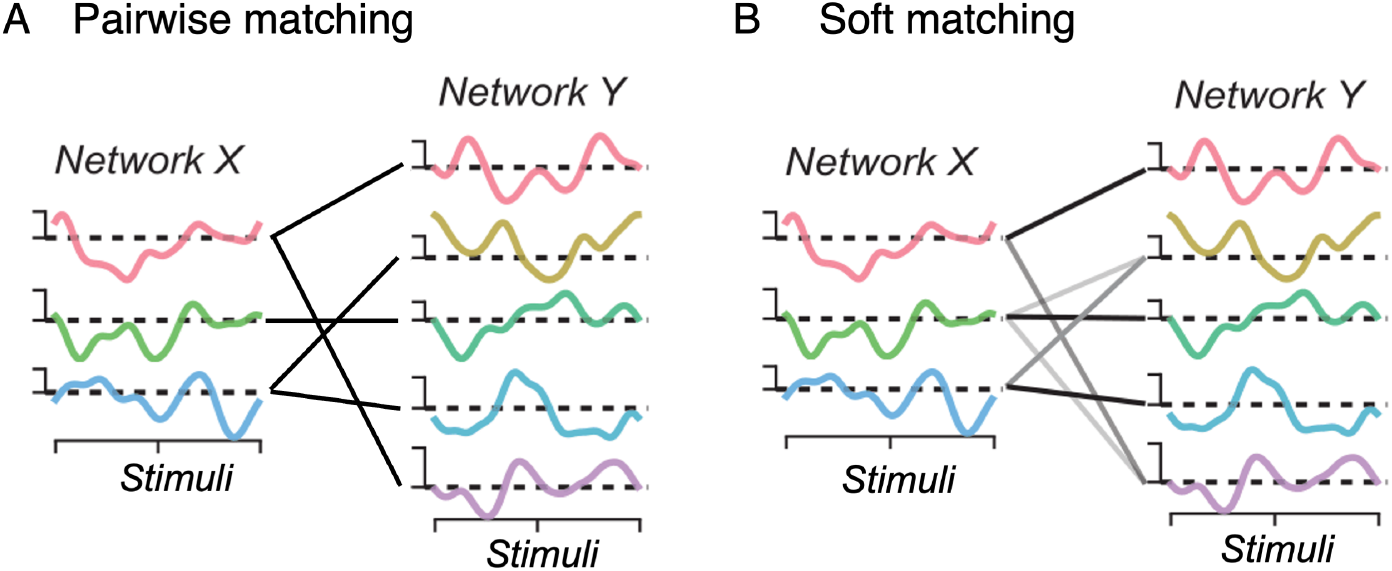
Example tuning curves from neurons in two representations over a 1D stimulus space. **A**Schematic illustration of pairwise matching. Lines show matched units across the two representations. **B** Schematic illustration of soft matching. Grayscale lines show matched similar tuning curves across two representations. The darkness of the line indicates the strength of the match.

Note that this measure is asymmetric by design. The choice of allowing repetitions with a bipartite semi-matching assignment (i.e. allowing a single unit in Y to be matched to multiple units in X) instead of a strict bipartite matching is motivated by potential variations in the number of neurons/units across different systems such as different artificial or biological networks, which may lead to varying levels of redundancy.

##### Soft Matching score

We also used a second similarity measure designed to address some limitations of the Pairwise Matching score, specifically that it is asymmetric (it yields different values depending on which of two representations is used as the “reference”) and does not satisfy the triangle inequality. This alternative measure satisfies all the requirements of a proper metric space.

Consider again two matrices 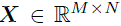 and 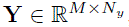, which represent the activations of *Nx* and *Ny* units respectively, for M instances. If *Nx* = *Ny* = *N*, instead of finding the optimal bipartite semi-matching assignment as in the direct pairwise matching score, we can attempt to find the optimal assignment between units without replacement, also known as a linear assignment problem [94] and as bipartite matching in graph theory [32]. This yields a *one-to-one* matching distance between representations,

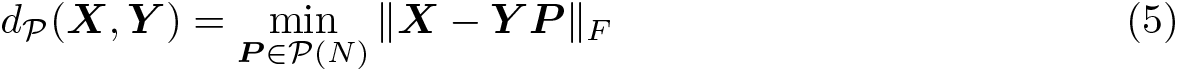

where the optimization is performed over the group of *N* –dimensional permutation matrices, *P*(*N*). The number of possible permutations is factorial in *N*, making brute force search impossible. However, an efficient algorithm can be achieved by replacing the space of permutations by the space of doubly stochastic matrices [95, 96]: square, nonnegative matrices whose rows and columns sum to one, which form the *Birkoff polytope B*(*N*). The resulting optimization problem is convex, and its solution can be shown to coincide with the optimal permutation.

Experimental datasets and neural network models typically have different numbers of neurons/units, meaning that no one-to-one matching of units is possible. Thus, we employ a generalization of the one-to-one matching as proposed in recent work [33]. To do this, we relax the set of permutations to “soft permutations” which result from extending the *Birkoff polytope* to rectangular matrices. Specifically, consider a nonnegative matrix 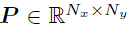 whose rows each sum to 1*/Nx* and whose columns each sum to 1*/Ny*. The set of all such matrices defines a *transportation polytope* [97], denoted as *T*(*Nx, Ny*). Optimizing over this set of rectangular matrices results in a “soft matching” or “soft permutation” of neuron labels in the sense that every row and column of ***P*** may have more than one non-zero element as shown in Figure 10B.

The expression for the *Soft Matching score* is:

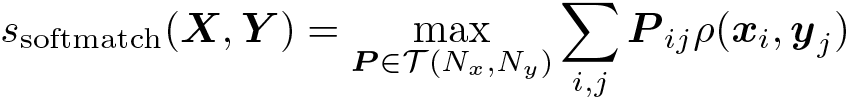

where *ρ*(***x****i, **y**j*) is the Pearson correlation between neuron *i* in network ***X***, and neuron *j* in network ***Y***. The score ranges between 0 and 1, providing a normalized alternative to *d*T. This measure has the appealing theoretical properties of optimal transport distances. Intuitively, the optimal mapping matrix ***P*** ∗ obtained by solving the optimization can be interpreted as a soft-permutation or soft-matching of neurons across the two representations.

Computing the above distance measures involves the resolution of a linear program, for which we employ efficient implementations of the simplex algorithm as in the Python Optimal Transport Library [98]. The calculation of the Soft Matching score was computationally expensive, and was prohibitive for the inter-subject comparisons of visual fMRI data, which had many voxels per subject. As a result, the Soft Matching score was omitted from analyses that involve intersubject comparisons for visual fMRI data. We included it in all other analyses to show that our conclusions generalize across two measures of axis alignment.

#### Quantifying alignment with a change of basis

We draw a random rotation matrix ***Q*** from the Haar distribution (the only uniform distribution on the special orthogonal group *SO*(*n*)) using an efficient algorithm based on *QR* decomposition implemented via Householder transformations [99]. The Haar measure ensures that each special orthogonal matrix is equally likely to be chosen.

We also consider smaller rotations (closer to the identity matrix ***I***) along the *SO*(*n*) manifold by computing fractional powers of the sampled rotation matrix, ***Q****α*(0 *< α <* 1). These fractional powers are computed by (i) triangulating ***Q*** into an upper quasi-triangular matrix ***T*** using the Schur decomposition (***Q*** = ***WT W*** ∗), (ii) computing ***T*** *α* by estimating the fractional powers of the individual 2 × 2 blocks comprising *T*, and (iii) reconstructing the fractional power of the original rotation matrix ***Q*** as ***Q****α* = ***WT*** *α**W*** ∗. For each representation pair (***X***, ***Y***), we change the basis of one representation, say ***X***, along this smooth SO(n) manifold to yield a transformed representation, ***X****′* = ***XQ****α*, and quantify how our axis-dependent alignment metric *s* (either *s*pairwise or *s*softmatch) changes with the amount of axis rotation (*α*) by computing *s*(***XQ****α, **Y***). If increasing rotations (from ***I*** to ***Q***) consistently reduce the alignment between representations, it provides empirical evidence that the system has a privileged basis. We note that each rotated representation has exactly the same linearly decodable information and population geometry and differs only in the tuning of its coordinate axes. In all experiments studying the influence of axis rotations on representational alignment, we stochastically sample ***Q*** 10 times and compute the alignment values across all sampled rotations for 10 values of *α* (spaced equally within the range [0.1, 1]).

##### Sampling rotation matrices as a composition of Plane (2D) rotations

In additional analyses, we investigated the effect of small axis perturbations on alignment between neural representations by sampling the special orthogonal (rotation) matrices using a different technique. Specifically, we constructed rotations in *n* dimensions as the product of *n*(*n* − 1)*/*2 plane rotations [100–102]:

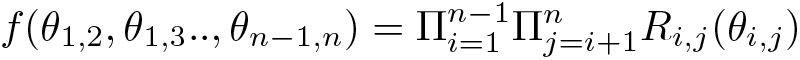

where *Ri,j* (*θi,j*) denotes the rotation in the *i, j* plane embedded in the *n*-dimensional representation with angle *θi,j*. Each 2D rotation affects only two coordinates at a time, leaving the others unchanged. For example, rotating a 3-dimensional representation can be achieved by combining individual rotations in each of the 2D planes within the 3-dimensional space. Parameterizing these single rotations with angles (*θ*1,2*, θ*1,3*, θ*2,3), we can, for instance, write *R*1,3(*θ*1,3) as,

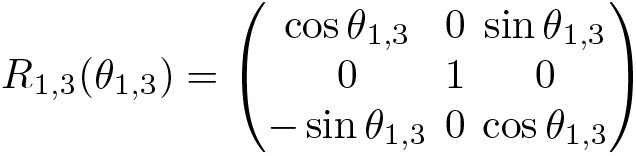

The combined rotation matrix can then be obtained by multiplying these matrices,

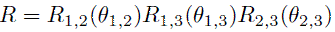

By sampling rotation matrices in this manner, we controlled *θi,j* to set the amount of each 2D rotation. We set *θi,j* to be a constant across all *i, j*, choosing from *π/*4*, π/*16*, π/*32*, π/*64. For each of these angles, we construct the final rotation matrices by composing a varying number (*c*) of 2D rotations, drawing *c* from 10 linearly spaced values between 0 and *n*(*n* − 1)*/*2. Here, *c* = *n*(*n* − 1)*/*2 corresponds to the rotation of all possible planes in an *n*-dimensional representational space.

We then applied these rotations to one representation within each pair of neural representations and studied how the angle of 2D rotation (*θi,j π/*4*, π/*16*, π/*32*, π/*64) and the number of 2D rotations (*c* ∈ [0*, n*(*n* − 1)*/*2]) affected alignment, as measured using pairwise matching. We performed this sensitivity analyses on brain-to-brain alignment as a proof of concept, and found that higher rotation angles (*θi,j*) and higher number of 2D rotations (*c*) more strongly reduced alignment. These results are presented in Supplementary Fig. S9.

#### Finding the optimal axis alignment

The procedure described in the previous section can reveal whether the alignment between axes in different instances of the system is higher in their native basis compared to a uniformly drawn basis. But such a result would leave open the possibility that the alignment is not as high as it could be, i.e., that there is a rotation matrix ***Q***∗ in the *SO(n)* manifold that yields a higher alignment than the standard basis. To assess whether the standard basis yields close to the maximum possible alignment, we used the Lie exponential and perform gradient-based optimization on the *SO(n)* matrix manifold [34] to find ***Q***∗ and compute this maximal alignment. The difference between the maximal basis alignment and natural basis alignment reveals the extent to which a representational system has a privileged basis.

More formally, we seek to find the optimal axis alignment between any pair of representations (***X****, **Y***). For an alignment metric *s* (either*spairwise* or *ssoftmatch*), we compute the optimal alignment *s*∗:

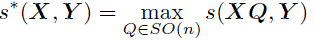

The above optimization is a constrained optimization problem. However, with a reparametrization, this constraint optimization can be turned into an unconstrained optimization, relying on the fact that every rotation matrix (***Q***) can be expressed as matrix exponential of a skew-symmetric matrix (***S***):

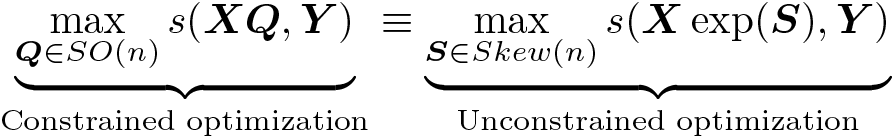

We found the solution for this unconstrained optimization problem using a regular gradient descent optimizer (Adam) by making use of the Geotorch library [34]. We used a learning rate of

0.001 and ran the optimization algorithm for 40 epochs in all experiments. Training curves were monitored for each experiment to ensure convergence.

### Analyses of the dominant components underlying DCNN representations

#### Bayesian non-negative matrix factorization algorithm

We identified the dominant components underlying DCNN representations using Bayesian nonnegative matrix factorization (NMF) in which the data matrix (units x stimuli) is modeled as the product of two lower rank matrices, both of which are constrained to be non-negative. The first matrix (henceforth referred to as the response profile matrix) encodes the response profiles of each of a set of components to all stimuli and the second matrix (henceforth referred to as the component-by-unit weight matrix) specifies the relative contribution of all units to each component.

Mathematically, the Bayesian NMF algorithm models the data matrix ***D*** as

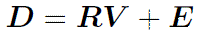

where ***D*** is the stimuli x units data matrix for a DCNN representation, ***R*** is the stimuli x components (N x C) response profile matrix, ***V*** is the components x units (***C*** x ***V***) weight matrix and ***E*** is a stimuli x units (N x V) residual matrix. In the Bayesian approach to NMF, ***R***, ***V***, and ***E*** are assumed to follow prior densities chosen for plausibility and mathematical convenience. For efficient inference, following Schmidt et al. [103], we chose a zero-mean normal residual matrix ***E*** with variance *σ*2, yielding a normal data likelihood:

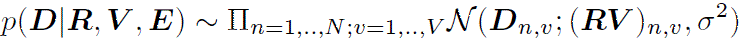

We further assume that ***R*** and ***V*** are independently exponentially distributed with scales *ρn,c* and *γc,v*:

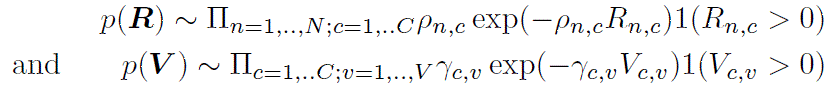

The conditional probabilities of ***R*** (i.e., *P* (***R D****, **V**, σ*2)) and ***V*** (i.e., *P* (***V D****, **R**, σ*2)) thus have a rectified Gaussian distribution. Following Schmidt et al. [103], the prior for the variance in ***E*** is assumed to have an inverse gamma distribution, resulting in an inverse-gamma conditional probability i.e., *P* (*σ*2 ***D****, **R**, **V***). Parameters for ***R***, ***V***, and ***E*** were optimized by sequentially drawing samples from these conditional densities using the Bayesian Markov Chain Monte Carlo (MCMC) sampling method derived in Schmidt et al. [103]. Due to the non-negativity constraints, the components identified by NMF are axis-dependent (in the sense that rotating the data will result in different components) unlike other methods like Principal Components Analysis (PCA).

#### Bayesian NMF Analysis of Visual Representations

We fix the number of components for representations in the self-supervised visual DCNN (’MoCo’) to 20, same as the optimal number of components estimated using the Bayesian Information Criterion (BIC) for the high-dimensional responses in the human ventral visual pathway in the NSD [19]. For obtaining the NMF components, we use the stimulus set shown to each of the 4 NSD participants who had full 3 repetitions for each of their 10,000 images. For the 10,000 images corresponding to each NSD subject’s data, we extracted the DCNN components separately. Further, since Bayesian NMF is a stochastic algorithm sensitive to initialization, we ran the algorithm N=10 times on the data matrix, each time obtaining C=20 components. We next compared these DCNN components with the 20 components obtained by applying the same factorization approach to high-level ventral visual stream responses in the four NSD participants. For each of these 4 × 10 different iterations of NMF yielding 20 components each, we identified the optimal assignment of those components to the five ventral visual stream components that were most consistent across NSD participants (see details in [19]). These components were previously shown to be selective to faces, scenes, bodies, text and food respectively [19]. Here, we first computed the mean response across participants of each of these five neural components to the 515 images viewed by all NSD participants. We next computed the negative pairwise correlation of the response of each of these five neural components with the response of the 20 DCNN components and applied the linear sum assignment algorithm [104] on the resulting correlation matrix to match each of the DCNN components to the most similar neural component. We report the mean correlations between the resulting 5 visual DCNN components and their corresponding neural counterparts across all 40 iterations normalized by the noise ceiling of each neural component (Figure 5B, left panel; top). The noise ceiling of each neural component was calculated by splitting the four NSD participants into two groups, computing the correlation between the mean response profiles of the component within each group, averaging the resulting correlation across all possible splits, and applying the Spearman-Brown correction.

For each of the DCNN components, we also report the log-kurtosis (a measure of sparsity) to reveal the extent to which component responses are driven by a few specific units as opposed to distributed responses among all units (Figure 5B, left panel; bottom). We also compute the correlation of each of the DCNN component responses with the salience ratings for their preferred category as obtained in an independent behavioral experiment [19] (reported in Figure 5B, middle panel insets).

#### Bayesian NMF Analysis of Auditory Representations

We determined the optimal number of components for representations in the self-supervised auditory DCNN (’BYOL-a’) to be 10. This was achieved by systematically varying the component count from 5 to 25 in intervals of 5, and determining the number that minimized the Bayesian Information Criterion (BIC). The NMF components were derived using model activations for the 165 sounds from the auditory fMRI dataset [37]. We again ran the Bayesian NMF algorithm N=10 times on the data matrix, each time obtaining C=10 components. For each of these 10 different iterations, we identified the optimal assignment of the 10 components to the two auditory fMRI components previously shown to be selective to speech and music [37, 105] using a linear assignment procedure analogous to that used for the visual NMF analysis. We report the mean correlations between the resulting 2 auditory DCNN components and their corresponding neural counterparts respectively across all 10 iterations (Figure 5C, left panel; top). We also computed the correlation of each of these DCNN component responses with a binary vector encoding the presence/absence of their preferred category of stimuli, i.e. ‘music’/‘speech’ (reported in Figure 5C, middle panel insets). We conducted additional control analyses to determine whether speech and music selectivity in DCNNs could be attributed to generic acoustic features, such as spectrotemporal modulations. For this purpose, we analyzed the component responses to a different set of 36 natural sounds and their modulation-matched counterparts, as used in Norman-Haignere et al. [105]. These component responses, derived from sounds different from the original 165 sounds in the study, were estimated by multiplying the DCNN activations by the pseudoinverse of the component weights.

### Analyses of category-selectivity in individual units of DCNNs and single voxels of fMRI data

The Bayesian NMF decomposition analysis reveals the dominant components underlying DCNN representations but does not tell us whether category-selective tuning is more prominent in the native basis of these representations than in arbitrary bases. To assess category-selectivity across different axes of visual DCNN and ventral visual stream representations, we used an image set encompassing characters, bodies, faces, and places [52] that is commonly used to localize category-selective regions in fMRI experiments (stimuli available at http://vpnl.stanford.edu/fLoc/). There were ten categories in this stimulus set, grouped into five stimulus domains: characters (word and number), bodies (body and limb), faces (adult and child), places (corridor and house) and objects (car and instrument). Stimuli consisted of an object from the category superimposed on a scrambled background (different backgrounds for different stimuli).

The same 10-category image set was also presented to NSD participants in separate functional localizer (fLoc) experimental runs with six repetitions of each stimulus. We used the GLM betas from the fLoc runs as the ventral visual stream representation in each NSD participant.

For each of the DCNN (MoCo) and cortical representations, we computed t-values to quantify the selectivity of individual units (single filters in DCNNs, single voxels in fMRI fLoC data) for different image categories. For instance, selectivity for faces was quantified as a t-value contrasting adult and child faces vs. all other categories. We then computed the number of units selective for each category using a t-value threshold of 10. We also assessed how the number of selective units changed under a shift of basis for each representation. As in our methodological framework for computing representational alignment, we drew a random rotation matrix ***Q*** from a uniform distribution on the special orthogonal group SO(n) to change the basis and considered smaller rotations along the SO(n) manifold by computing fractional powers of the sampled rotation matrix, ***Q****α*, where *α* ranged across 10 equally spaced intervals within [0.1, 1]. We stochastically sampled rotation matrices 10 times and computed the mean number of selective units across all sampled rotations for each *α*.

In the auditory domain, we used the 165 sounds used to obtain the fMRI dataset to identify category-selective units (we did not have data from any independent functional localizer runs in the auditory domain). For both the auditory cortex and auditory DCNN (BYOL-a) representations, we computed the number of units selective for music and speech separately using a t-value threshold of 10. Here, t-values were computed by contrasting the response to each category of sound (music or speech) vs. all other sounds. We followed the same procedure as in the visual domain for assessing how the number of selective units changes with an altered basis.

### Comparison of DCNNs and transformer architectures with different metrics

We compared the representation of each DCNN/Transformer model to that of the corresponding sensory areas of the human brain using the two axis-dependent alignment measures described above (pairwise matching and soft matching scores) as well as the arguably most widely used brain-model similarity metric (linear predictivity). For computing linear predictivity scores, we followed the standard procedure for fitting task-optimized models to brain responses based on a full linear readout. We extracted features from penultimate pre-trained layers of each network and fitted *l*2 regularized linear regression models separately for each network. The regularization parameter was selected by first fitting a regression model for each parameter among 8 log-spaced values in [1e-4, 1e4], using a training split of data, and then selecting the parameter value that gave the best accuracy on a validation split. This selection was performed independently for each animal/subject and each neural network model. We quantified ‘linear predictive accuracy’ on the test stimuli as the Pearson correlation coefficient between the predicted and measured response at each voxel or neuron. Here, we use the same train-val-test split as used for the pairwise matching correlation scores and described under the respective section for each dataset.

Because the human visual fMRI data were derived from an event-related design, the voxel responses were relatively noisy, and so we noise-corrected the linear predictivity estimates to give a better sense of the quality of the model predictions (Figure 7A). We performed the correction using the noise ceiling for each voxel, which were computed using the standard procedure followed in [35] by considering the variability in voxel responses across repeat scans. We also noise-corrected the Pairwise Matching score, as this was natural given that the score is based on a correlation. Because there was not a natural way to noise-correct the Soft Matching scores, we report the raw scores for this measure. The auditory fMRI data were derived from a block design in which each sound was presented multiple times in a row, and were substantially more reliable than the visual data, and so were not noise-corrected.

Following the idea that a good model should be similar to the brain in the same sense that conspecifics are similar to each other [106], we also compared the values of our axis-alignment measure (Pairwise Matching score) to an ‘inter-animal’ or ‘inter-human’ consistency score derived for each dataset. This consistency score was derived using an identical train-val-test split as that described above. The consistency score was averaged across all pairs of humans/animals.

### Analyses of computational implications of privileged axes

#### Efficiency

We operationalized efficiency in terms of how sparsely a unit/neuron responds to all presented natural stimuli (’lifetime sparseness’). We quantified sparseness as the Fisher’s kurtosis (*κ*4) of the response distributions of each unit/neuron in response to all presented stimuli (a normal distribution of such responses across stimuli would yield *κ*4 = 0). This lifetime sparseness measure was calculated for individual units/voxels/neurons and then averaged across all units/voxels/neurons. We assessed how sparseness changes with axis perturbations by again drawing a random rotation matrix ***Q*** from a uniform distribution on the special orthogonal group *SO(n)* to change the basis and considered smaller rotations along the *SO(n)* manifold by computing fractional powers of the sampled rotation matrix, ***Q****α*, where *α* ranged across 10 equally spaced intervals within [0.1, 1]. We stochastically sampled these rotation matrices 10 times and compute the mean sparseness values across all sampled rotations for each *α*. We applied this same procedure to single-neuron recordings in macaque IT (as in the Macaque-NSD dataset), intracranial recordings in human auditory cortex [48], and penultimate layer activations in self-supervised visual (‘MoCo’) and auditory (‘BYOL-a’) networks.

Since kurtosis could distort sparsity comparisons due to the nonnegative response distributions in the native axes, we also employed another measure of lifetime sparseness previously proposed by Vinje and Gallant [58]. Here, sparseness (S) for a set of responses {*ri*}*i* =1*,..,n* where *ri* denotes the response of the neuron/unit to the *i*th stimulus and *n* denotes the number of stimuli is quantified as

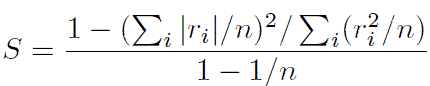

Results obtained using this metric are reported in Supplementary Fig. S12.

#### Downstream wiring costs

We also tested whether the axes found in the brain are those that facilitate a sparse readout of behaviorally relevant information (e.g. object category labels) from patterns of neural activity. To explore this hypothesis, we employed linear decoders trained on neural representations from both computational models and actual brain data, tasked with predicting stimulus categories of interest. In the visual domain, we used the shared 1,000 NSD images for extracting both model (here, the self-supervised visual DCNN trained using momentum contrast) and brain representations, which included human ventral visual stream responses in the NSD and macaque IT responses in the Macaque-NSD dataset. The decoder classes were the 80 object categories in the MS-COCO dataset from which the NSD stimuli were drawn. Since an image in the NSD can have more than one object category, we set this up as a multi-label classification task. We trained separate ridge classification models on each visual representation. The model was regularized with an *l*2-penalty, with the aim of ensuring equivalent performance across all representational axes (since changing the axes does not alter the *l*2-norm). The regularization parameter was selected by first fitting a regression model for each parameter among 8 log-spaced values in [1e-4, 1e4], using a training split of data, and then selecting the parameter value that gave the best accuracy on a validation split. This selection was performed independently for each brain and model representation. We quantified the classification performance for each category label separately with the Area under the Receiver Operating Characteristic (AUC-ROC) measure and computed the mean across all categories to derive the aggregate prediction performance. We estimated the final prediction performance by averaging the results across five splits of the data for each representation, holding out one-third of the data for testing in each split. We computed the readout sparsity by estimating the sparseness of the readout weight distribution of each category over all units/neurons/voxels using Fisher’s kurtosis (*κ*4). Aggregate readout sparsity was computed by taking the mean across all categories (Figure 8B, top right panels)).

We again repeated the procedure for different axis perturbations by drawing random rotation matrices ***Q*** from a uniform distribution on the special orthogonal group *SO(n)* to change the basis and considered smaller rotations along the *SO(n)* manifold by computing fractional powers of the sampled rotation matrices, ***Q****α*, where *α* ranged across 10 equally spaced intervals within [0.1, 1].

We stochastically sampled these rotation matrices 10 times and computed the mean classification performance and readout sparsity across all sampled rotations for each *α*. We also visualized the difference between the readout sparsity in the native axes and mean sparsity over all fully-rotated axes (*α* = 1) for each of the 80 categories as a histogram (Figure 8B, bottom right panels).

In the auditory domain, we followed a similar procedure as in the visual domain, training ridge classification models on intracranial cortical responses and self-supervised DCNN (‘BYOL-a’) activations to natural auditory sounds. The classifier decoded the category labels of the sounds, which spanned eleven human-interpretable categories (environmental sounds, English speech, foreign speech, animal vocalizations, animal non-vocalizations, human vocalizations, human non-vocalizations, mechanical sounds, music, nature sounds and song). The regularization strength parameter was optimized independently for each representation by validating among 9 log-spaced values in [1e-4, 1e4]. We evaluated the classification performance by computing the mean AUC-ROC measure (one vs. rest) using the predicted probability of each stimulus category. We estimated the classification performance by averaging results across five splits of the data, holding out one-third of the data for testing in each split, as in the visual domain. We computed the readout sparsity by estimating the sparseness of the readout weight distribution of each category over all units/electrodes using Fisher’s kurtosis (*κ*4). We followed the same approach for comparison against arbitrary uniformly-drawn bases as described for the visual representations.

#### Few-shot generalization

We also tested the hypothesis that privileged axes could improve generalization for certain tasks under constrained learning scenarios. Specifically, we hypothesized that privileged axes would result in improved generalization capabilities for predicting categories whose selective responses are manifested in these axes, and that these benefits would be most evident during constrained learning scenarios such as few-shot learning or learning with enforced sparsity on the downstream linear readout. This hypothesis is based on a theoretical assertion described in the next section. To test this hypothesis empirically, we trained sparse linear decoders on top of neural representations in both native and arbitrary bases. A sparsity constraint, meant to mimic anatomical constraints on the number of inter-region connections, was enforced on the decoder through an *l*1 penalty. The *l*1 penalty also makes the generalization performance of the decoder axis-dependent. A few-shot learning scenario was instantiated by training the linear decoders using only a small fraction (10%) of labeled stimulus-category pairs. In the visual domain, separate sparse linear regressors (Lasso) were trained to predict the presence of each of five categories in an image: faces, places, bodies, text and food. The presence of each category was quantified using human salience ratings. These salience ratings were collected in an independent behavioral experiment [19] by asking participants how noticeable each of the five categories was in every image. For auditory representations, two distinct sparse linear classifiers (Logistic regression with *l*1 penalty) were trained to detect the presence of music or speech in a sound waveform. The regularization parameter for the *l*1 penalty was selected by first fitting a regression model for each parameter among 8 log-spaced values in [1e-4, 1e4], using a training split of data, and then selecting the parameter value that gave the best accuracy on a validation split. This selection was performed independently for each task and for each model and brain representation. We considered representations from the human ventral visual stream and the self-supervised visual DCNN (‘MoCo’) in response to all images for which salience ratings were available (515 in total). In the auditory domain, we considered representations from the human auditory cortex as measured using intracranial recordings and the self-supervised auditory DCNN (‘BYOL-a’) in response to all 165 sounds. We performed comparisons against arbitrary axes by again drawing random rotation matrices ***Q*** from a uniform distribution on the special orthogonal group *SO(n)* to change the basis. We stochastically sampled these rotation matrices 10 times and computed the mean value of the classification metrics across all sampled rotations.

For each candidate representation and basis (native vs. arbitrary), we computed the generalization performance of the trained sparse linear decoders on held-out stimuli. For the visual tasks of predicting category salience, we quantified the generalization performance by computing the coefficient of determination *R*2 between the predicted and measured responses. For the auditory tasks of predicting category presence vs. absence, we quantified generalization performance by computing the AUC-ROC by using the predicted probabilities for each class. We aggregated the performance metrics across 5 splits of the data using a 10 − 90 train-test split corresponding to the few-shot scenario.

##### Why do privileged axes facilitate few-shot learning?

Here we provide a mathematical explanation for why certain axes facilitate few-shot learning when using sparsity constraints on the downstream readout. Consider the representation 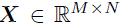 as a matrix encoding the activations of *N* neurons for *M* stimuli. Let 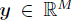 be a target variable that represents a meaningful factor of variation in our external world. We assume that it is continuous-valued, such as the salience of a behaviorally-relevant category in a stimulus. The readout neuron learns to encode this factor from these *M* instances of stimuli. Here, we focus on the learning problem where the representation is fixed and only the readout weights ***w*** ∈ ℝ^*N*^ are learned. Assuming a linear readout, the weights can be learned within a linear estimation framework:

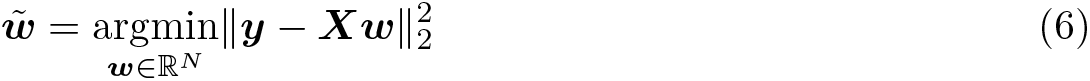

In many such situations in biological nervous systems, the readout neuron may not have access to all the neurons of a representation due to wiring constraints. Thus, ***w*** 0 = *N* ′ for some *N* ′ *< N*. During learning, this assumption can be instantiated as a sparsity constraint on the learned weights ∥***w***∥0. Under this assumption, the learning problem can be expressed as follows:

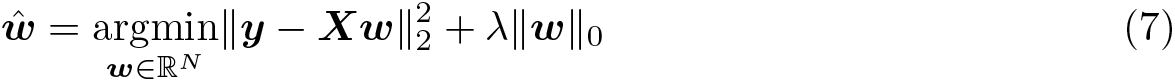

where *λ* is a regularization parameter that controls the strength of the sparsity penalty. The generalization error of the above estimation problem can be decomposed into bias and variance terms. The expected error can be expressed in terms of the ideal weights ***w***∗as

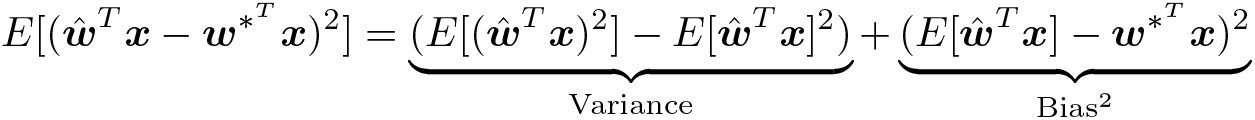

Although there is no closed-form solution for the bias and variance of the estimator subject to a sparsity constraint (Eq. 7), the bias-variance tradeoff suggests that reducing the size of the hypothesis class using sparsity-based regularization should reduce the variance of the estimator [107]. This would hold true for any representation ***X***, irrespective of the representational form it uses to encode the target variable ***y***. Consider a transformed representation ***X***′ = ***XR*** with equivalent representational content as ***X*** obtained by randomly rotating the axes of the representation ***X*** through multiplication with a rotation matrix ***R***. The solution to the unconstrained optimization problem ***w***′ = argmin***w***∈ ℝ^*N*^ ∥***y*** − ***X***′***w***∥2 can be expressed in terms of the solution ***w***~ to Eq. 6 as it simply reflects a reparametrization,

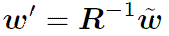

Now, let us assume that the standard basis representation ***X*** employs a sparse code to encode the target variable ***y***, i.e. ∥***w***~ ∥0 *< N*. In this case, the sparsity constraint in Eq. 7 does not

exclude the estimator ***w***~ from the hypothesis class, leaving the bias unaffected. On the other

hand, multiplication with a random ***R***−1 will generally destroy the sparsity of ***w***~, and therefore ***w***′ may no longer be included in the hypothesis space of Eq. 7. This would cause an increase in the bias term for the rotated representation ***X***′, increasing the overall generalization error of the estimator. Thus, generally, while the sparsity constraint would reduce the variance of the estimator for both ***X*** and ***X***′, it can increase the bias term for ***X***′resulting in worse generalization in comparison to ***X***. In practice, for all our experiments, we considered a convex relaxation of the learning problem described above 7 using the L1 norm instead of L0 norm since the convex relaxation can be easily solved using coordinate descent.

## Supplementary Information

### Simulations and Control Experiments

#### Proposed rotation-sensitive measures are sensitive to both representational content and form

We first present simulations demonstrating that the proposed pairwise matching and soft matching similarity scores effectively discriminate between representations that differ in the tuning of individual neurons. Consider one representation 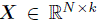 formed from sampling k linearly independent latent factors 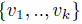 drawn from a normal distribution, 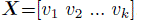 can be viewed as the activation space of *k* units for *N* stimuli. Now, consider a second representation ***Y*** derived as a linear combination of some of these factors using a mixing matrix 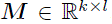: 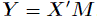, where 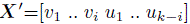 and 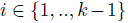. Here, the remaining latent factors {*u*1*, …, uk*−*i* } are independently drawn. The fraction *i/k* governs the alignment of latent factors between ***X*** and ***Y***: higher values result in increased alignment. Furthermore, the sparsity of ***M***, measured by the number of non-zero entries in ***M***, nnz(***M***), controls the similarity between the single-neuron tuning functions of ***X*** and ***Y***: a sparse ***M*** leads to an increased alignment of tuning functions. We varied the alignment of latent factors (representational content) and the alignment of tuning functions (representational form) and studied the sensitivity of different measures of representational similarity to these two types of alignment. Our empirical simulations, shown in Figure S1, confirm that the pairwise matching and the soft matching scores accurately capture stricter notions of representational similarity, extending beyond information content to encompass the alignment of single-neuron tuning functions as well. In contrast, conventional measures such as centered kernel alignment (CKA), representational similarity analysis (RSA) or linear predictivity are only sensitive to the alignment of latent factors (i.e. the information content), regardless of whether these factors are disentangled in one representation (here, ***X***) and entangled in the other (***Y***).

**Fig. S1.**
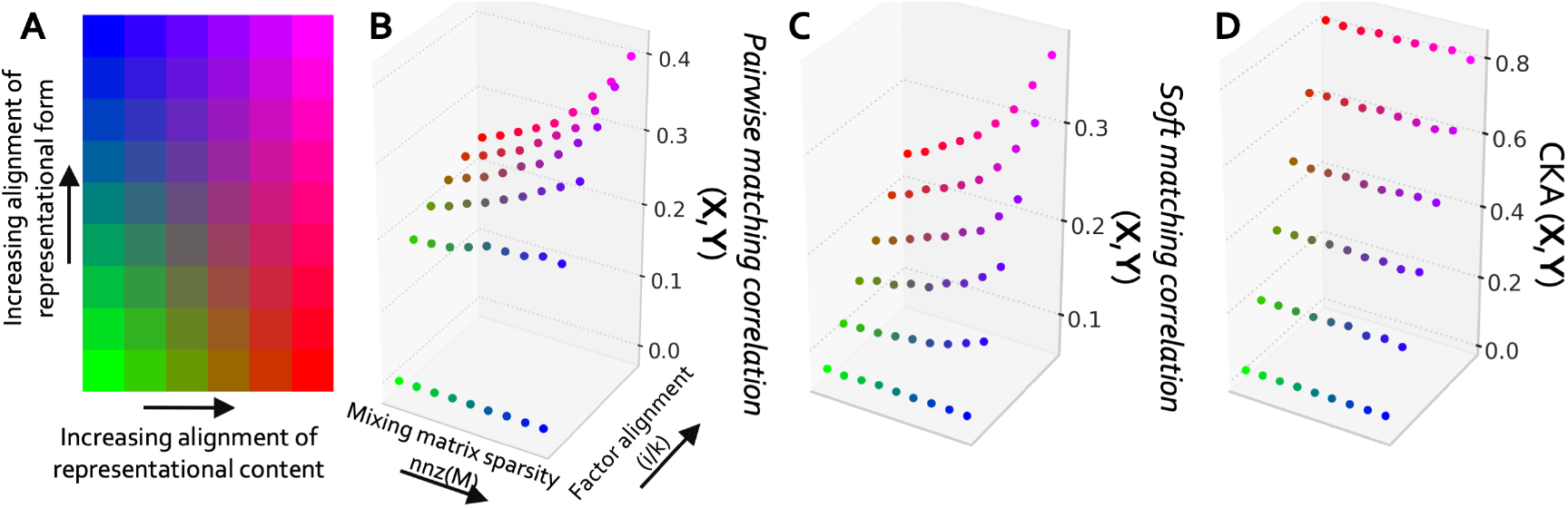
Proposed measures effectively capture neural tuning alignment. We sample two representations, ***X*** and ***Y***, varying the extent of alignment in their representational content and form, with the color scheme illustrated in (**A**). (**B**), (**C**) and (**D**) visualize the changes in alignment between ***X*** and ***Y*** with respect to these manipulated variables, as assessed using our pairwise matching score (*s*pairwise) soft matching score (*s*softmatch) and centered kernel alignment (CKA), respectively. CKA distinguishes representations that differ in content but is blind to differences in representational form, whereas our two new measures are sensitive to differences in both content and form. In this simulation, *N* = 4000, *k* = 100 and *l* = 1000.

#### Measuring alignment change with axis rotation reliably reveals the existence of privileged axes

We next present simulation results demonstrating that the proposed method of quantifying alignment change with axis rotations is diagnostic of the presence of a privileged axis within a representational system. We generated synthetic data varying in the presence of privileged axes, by mixing a set of factors using mixing coefficients that varied in sparsity. We first generated a matrix 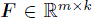 containing responses of *k* linearly independent factors to *m* stimulus conditions. Each column of ***F*** is sampled from a standard normal distribution, with the columns orthogonalized by the Gram-Schmidt procedure. We then generated two “neural” representations 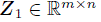 and 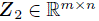 as linear mixtures of these factors:

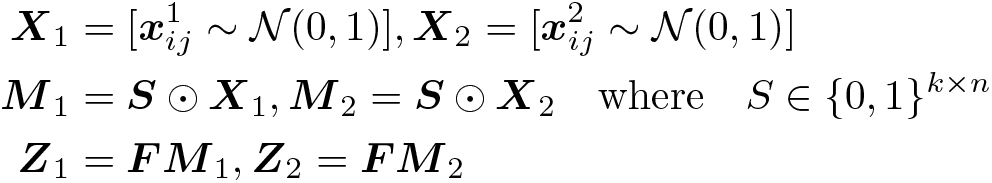

This way of generating ***Z***1 and ***Z***2 causes individual neurons to be mixtures of a subset of the factors in ***F***. The level of sparsity in the mixing matrices is controlled by the number of non-zero entries in ***S***. When ***S*** is highly sparse, ***Z***1 and ***Z***2 have privileged representational axes, with the “neurons” tending to be aligned with individual factors or small mixtures of factors in ***F***. When ***S*** is dense, each “neuron” in ***Z***1 and ***Z***2 is a linear mixture of all independent factors. In this case, there are not privileged axes, i.e. the two representations are no more aligned in their native axis than an arbitrary axis. Here, we empirically examine how the alignment between ***Z***1 and ***Z***2, determined using the proposed rotation-sensitive measures, varies as we gradually rotate the representational axis in either ***Z***1 or ***Z***2. We emphasize that standard measures, like centered kernel alignment (CKA), representational similarity analysis (RSA) or linear predictivity would not show any change with axis rotations. As shown in Figure S2, when ***S*** is sparse, i.e., when the representations have privileged axes, the alignment between ***Z***1 and ***Z***2 drops sharply with axis rotation. Conversely, when ***S*** is dense and the representations lack distinct axes, this trend is not observed. This observation confirms the effectiveness of our method in identifying the prevalence of a distinct or ‘privileged’ basis.

#### Changes in alignment with axis rotations are not driven by the changing statistical structure of neural/unit responses

In our proposed framework, we assess changes in alignment through axis rotations to determine if a representational system has a privileged axis. If axis rotations (sampled uniformly from the SO(n) group) consistently reduce the alignment between different instances of representations from a given system, it suggests the presence of a preferred axis. However, an alternative explanation could be that axis rotations alter the statistical distributions of unit/neuron activations, causing the observed alignment loss. For example, as demonstrated in Fig. 8A, the response distributions of units in their native axes was sparser (heavier-tailed and right-skewed) than that for units in an arbitrary basis. This pattern was consistent across both artificial and biological representations in vision and audition.

**Fig. S2.**
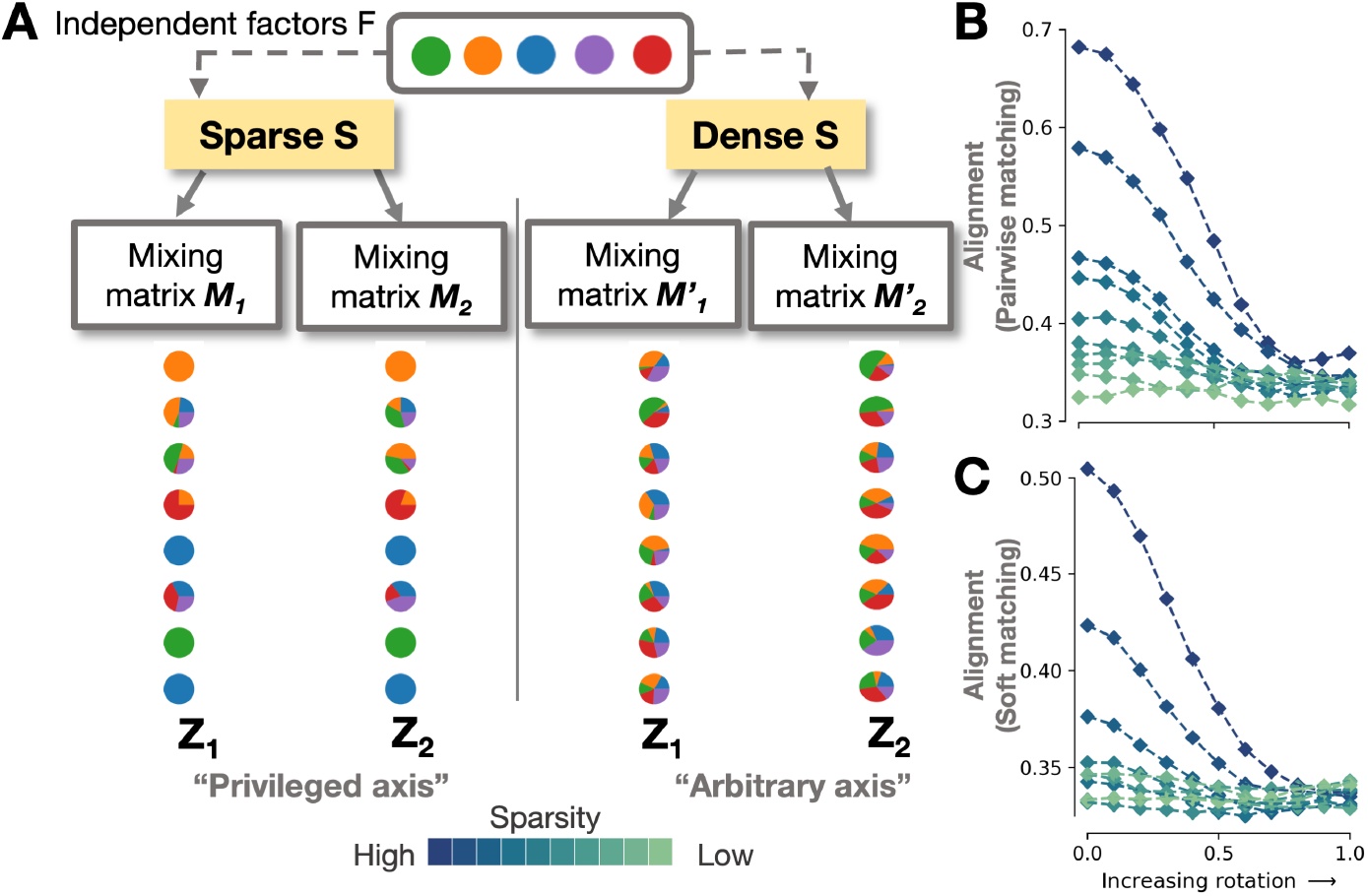
Investigating privileged axes through gradual axis rotations. (**A**) Illustrated schematic featuring two instances (***Z***1 and ***Z***2) of a representational system where each unit is a linear mixture of several independent latent factors. Colors depict the independent factors that each unit encodes. The sparsity of the mixing matrix controls the distinctiveness of the coordinate axis: high sparsity reflects the privileged axis scenario with a direct alignment of neural tuning functions, whereas low sparsity reflects the arbitrary axis scenario. (**B**) and (**C**) depict the change in alignment as a function of axis rotation (*α*), as measured using the pairwise matching score (*s*pairwise) and the soft matching correlation score (*s*softmatch), respectively. The dark blue curves correspond to the privileged axis scenarios, consistently displaying a gradual alignment decrease with axis rotations, in contrast to the light blue curves representing the arbitrary axis scenarios. While our illustration shows privileged axes aligning with independent data generative factors, the concept of privileged axes in our study is broader, and includes any consistent tuning function alignment across different system instances regardless of what the tuning functions encode. In this simulation, *m* = 1000, *k* = 50 and *n* = 100.

To test whether the differing response statistics can explain the alignment drop between native and arbitrary axes, we first conducted a simulation. We constructed two representations ***X*** and ***Y*** with unit responses drawn from a heavy-tailed right-skewed distribution, similar to the distributions observed for neural responses in their native axes (Fig. 8A). Specifically, the responses were sampled from a gamma distribution with a shape parameter (*k*) of 10 and a scale parameter (*θ*) of 1 (Fig. S3A). Since the responses were drawn independently for the two representation instances (***X*** and ***Y***), there should be no privileged axes shared between the two representations. If a heavy-tailed right-skewed distribution were sufficient to cause alignment to change with axis perturbations, one should expect the alignment between ***X*** and ***Y***, as measured using our axis-sensitive measures (pairwise matching and soft matching correlation scores), to drop with axis rotations (since axis rotations alter the response statistics). However, as demonstrated in Fig. S3A, even full axis rotations (*alpha* = 0) result in a non-significant change in the alignment values using either measure. This demonstrates that the changes in the distribution of unit/neuron responses after axis rotations does not explain the drop in alignment values. We note that for this experiment, when computing the pairwise matching score, we evaluated the correlation score on the same data that was used for finding the pairwise mappings (i.e. without cross-validation). Cross-validation would have resulted in scores hovering around zero due to the lack of a common stimulus dimension.

We conducted another control experiment to further demonstrate that the loss in alignment with increasing axis rotations can not be merely explained by the changing statistical structural of neural responses with axis rotations. We computed the alignment between neural responses in macaque IT (from the ManyMonkeys dataset) using the same brain-to-brain alignment procedure described in the main text. However, this time, we shuffled the stimulus ordering independently for each monkey. This shuffling preserves the response statistics of individual neurons while disrupting any common stimulus-driven structure.

**Fig. S3.**
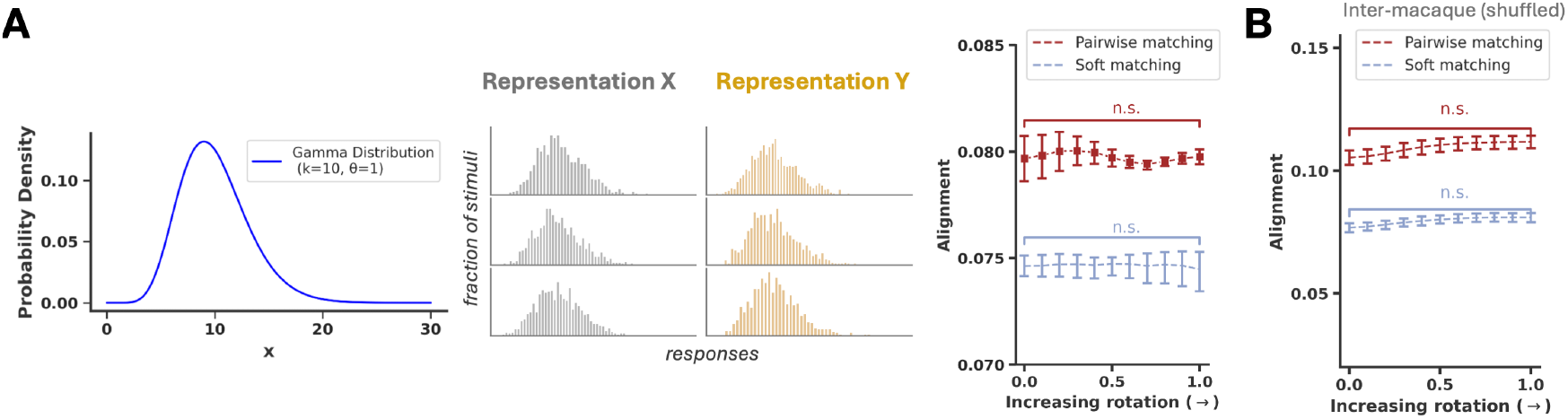
Marginal response distributions do not account for axis alignment. (**A**) Left to right: Density function of a gamma distribution with a shape parameter of 10 and a scale parameter of 1; response distributions of sample units in simulated representations ***X*** and ***Y*** sampled from the previous gamma distribution; Alignment between ***X*** and ***Y*** plotted against axis rotation, assessed using pairwise matching and soft matching correlation scores. (**B**) Alignment between conspecific neural responses in macaque IT after shuffling of the stimulus order (preserving the response distributions but eliminating any shared representational axes), evaluated with pairwise matching and soft matching correlation scores, and examined as a function of axis rotation (*α*).

We again observed that even full axis rotations (*α* = 0) resulted in no significant change in alignment between the shuffled neural representations in different monkeys, using both the pairwise matching and soft matching measures (Fig. S3B). This finding reinforces the conclusion that changes in alignment values with axis rotations are not driven by changes in the statistical structure of neural responses alone. As in the previous experiment, we did not perform crossvalidation when computing the pairwise matching score to avoid scores hovering around zero due to the lack of a common stimulus dimension.

## Additional Supplementary Figures

**Fig. S4.**
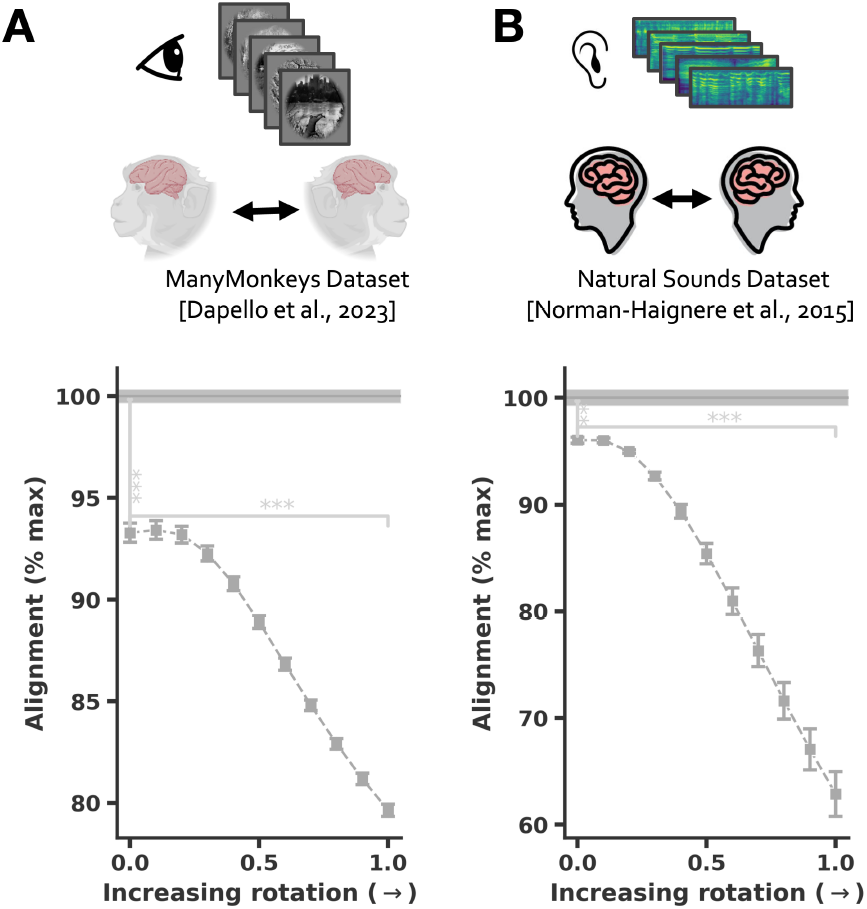
Brain-to-brain alignment using the soft matching score. (**A**) and (**B**) depict alignment between conspecifics neural responses in macaque IT and human auditory cortex. This alignment is measured using the soft matching score and examined as a function of axis rotation (*α*) where *α* = 0 indicates no rotation and *α* = 1 indicates full rotation using a rotation matrix drawn uniformly from SO(n). Alignment is expressed as a proportion of the maximum possible value obtainable across all axis rotations, determined using the ‘Optimal Axis Alignment’ procedure described in the text. Error bars are SEM over multiple splits of the data. Significance tests compare the alignment values at no rotation (*α* = 0) with those at full rotation (*α* = 1), as well as with the optimal alignment values observed across all rotation angles.

**Fig. S5.**
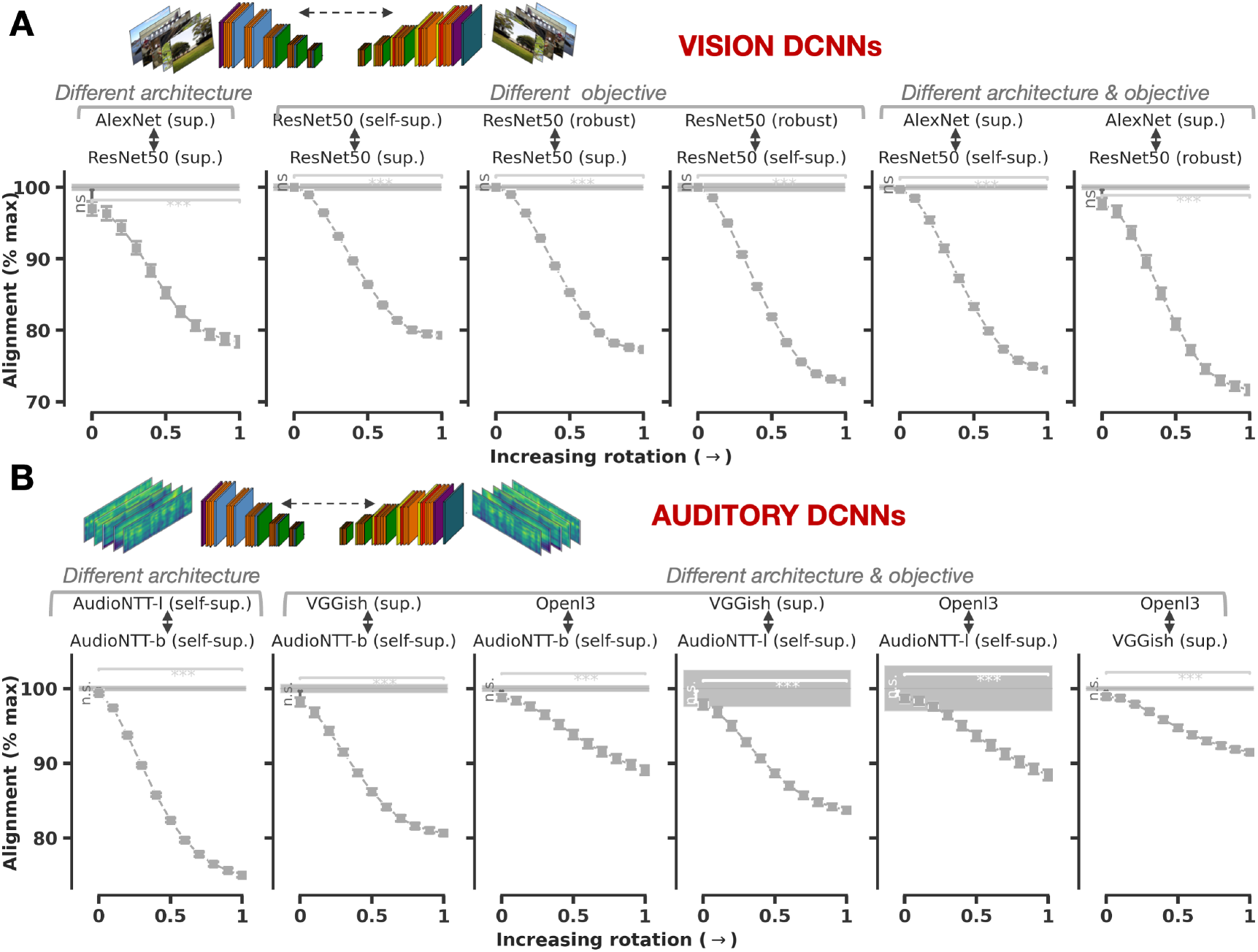
Model-to-Model alignment using the soft matching score. (**A**) Alignment between different visual deep convolutional neural networks trained with different architectures and learning objectives. This alignment is measured using the soft matching score and examined as a function of axis rotation (*α*) where *α* = 0 indicates no rotation and *α* = 1 indicates full rotation using a rotation matrix drawn uniformly from SO(n). Alignment is expressed as a proportion of the maximum possible value obtainable across all axis rotations, determined using the ‘Optimal Axis Alignment’ procedure described in the text. Alignment computed from activations to the 1000-image NSD stimulus set used for computing inter-subject alignment. (**B**) Same as A, but for auditory neural networks, and computed from activations to the 165 natural sounds used for auditory inter-subject alignment analysis. In each case, we gradually rotate the axis in one representation while computing its alignment with the other representation in its native basis. Error bars are SEM over multiple splits of the data. Significance tests compare the alignment values at no rotation (*α* = 0) with those at full rotation (*α* = 1), as well as with the optimal alignment values observed across all rotation angles.

**Fig. S6.**
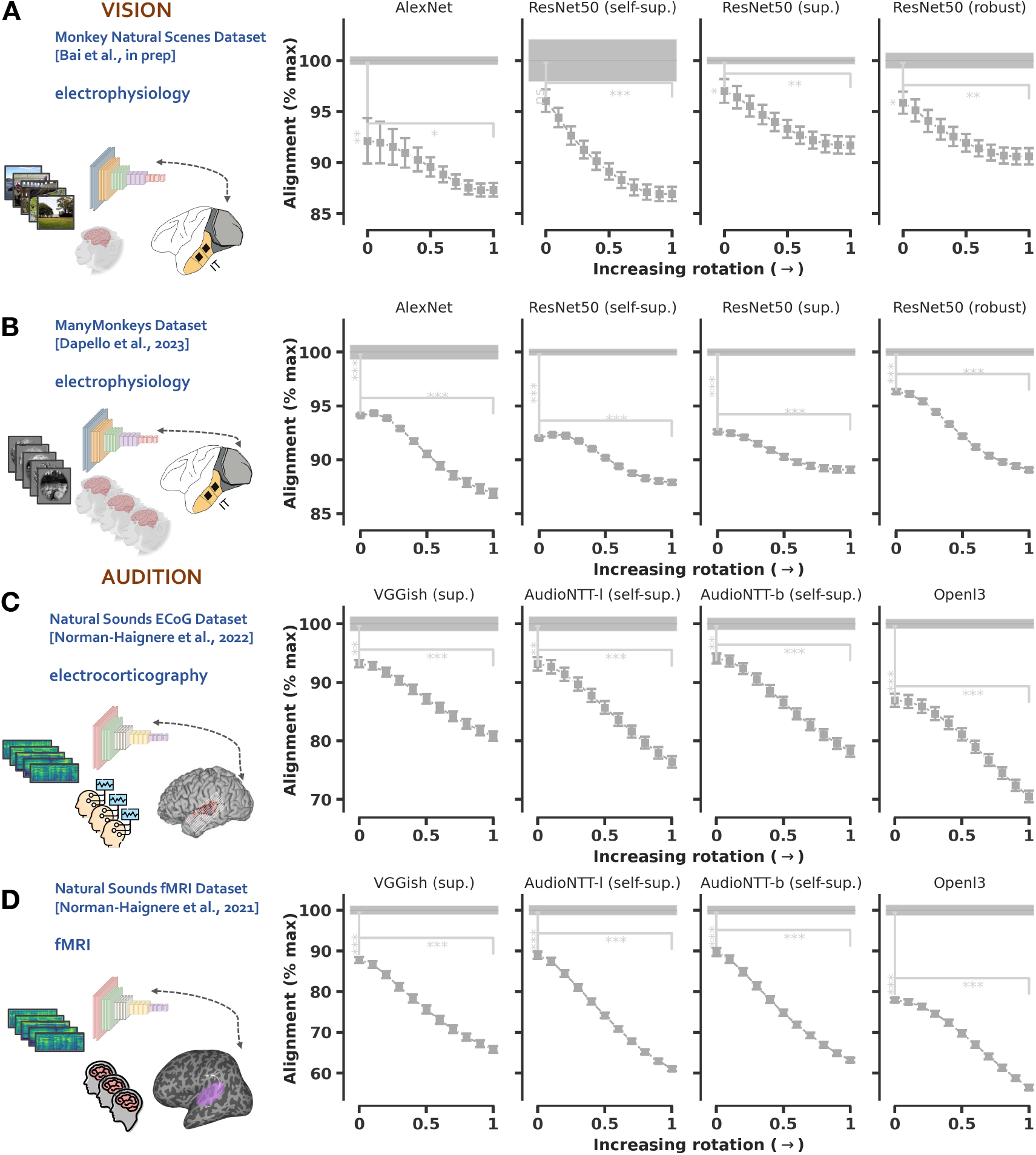
Model-to-brain alignment using the soft matching score. (**A-B**) Alignment of various visual deep convolutional neural networks with biological neural representations in the macaque IT. This alignment is measured using the soft matching correlation score and examined as a function of axis rotation (*α*) where *α* = 0 indicates no rotation and *α* = 1 indicates full rotation using a rotation matrix drawn uniformly from SO(n). Alignment is expressed as a proportion of the maximum possible value obtainable across all axis rotations, determined using the ‘Optimal Axis Alignment’ procedure described in the text. (**C-D**) Alignment of auditory DCNNs with neural representations in the human auditory cortex, again assessed as a function of DCNN axis rotation. Error bars are SEM over multiple splits of the data. Significance tests compare the alignment values at no rotation (*α* = 0) with those at full rotation (*α* = 1), as well as with the optimal alignment values observed across all rotation angles.

**Fig. S7.**
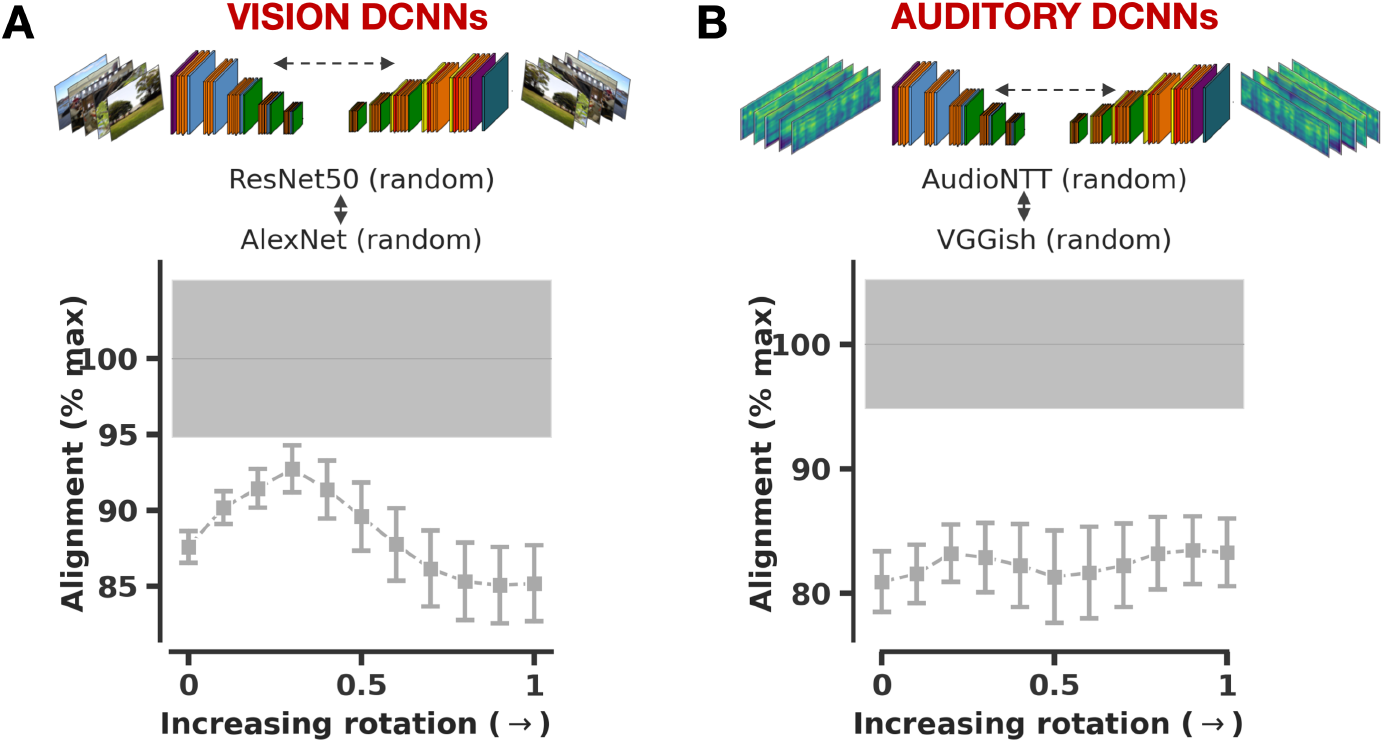
Model-to-Model alignment in untrained networks. (**A**) Alignment between untrained visual deep convolutional neural networks trained with different architectures (ResNet50 and AlexNet). This alignment is measured using the pairwise matching score and examined as a function of axis rotation (*α*) where *α* = 0 indicates no rotation and *α* = 1 indicates full rotation using a rotation matrix drawn uniformly from SO(n). Alignment is expressed as a proportion of the maximum possible value obtainable across all axis rotations, determined using the ‘Optimal Axis Alignment’ procedure described in the text. Alignment computed from activations to the 1000-image NSD stimulus set used for computing inter-subject alignment. (**B**) Same as A, but for untrained auditory neural networks with different architectures (AudioNTT and VGGish), and computed from activations to the 165 natural sounds used for auditory inter-subject alignment analysis. In each case, we gradually rotate the axis in one representation while computing its alignment with the other representation in its native basis. Error bars are SEM over multiple splits of the data.

**Fig. S8.**
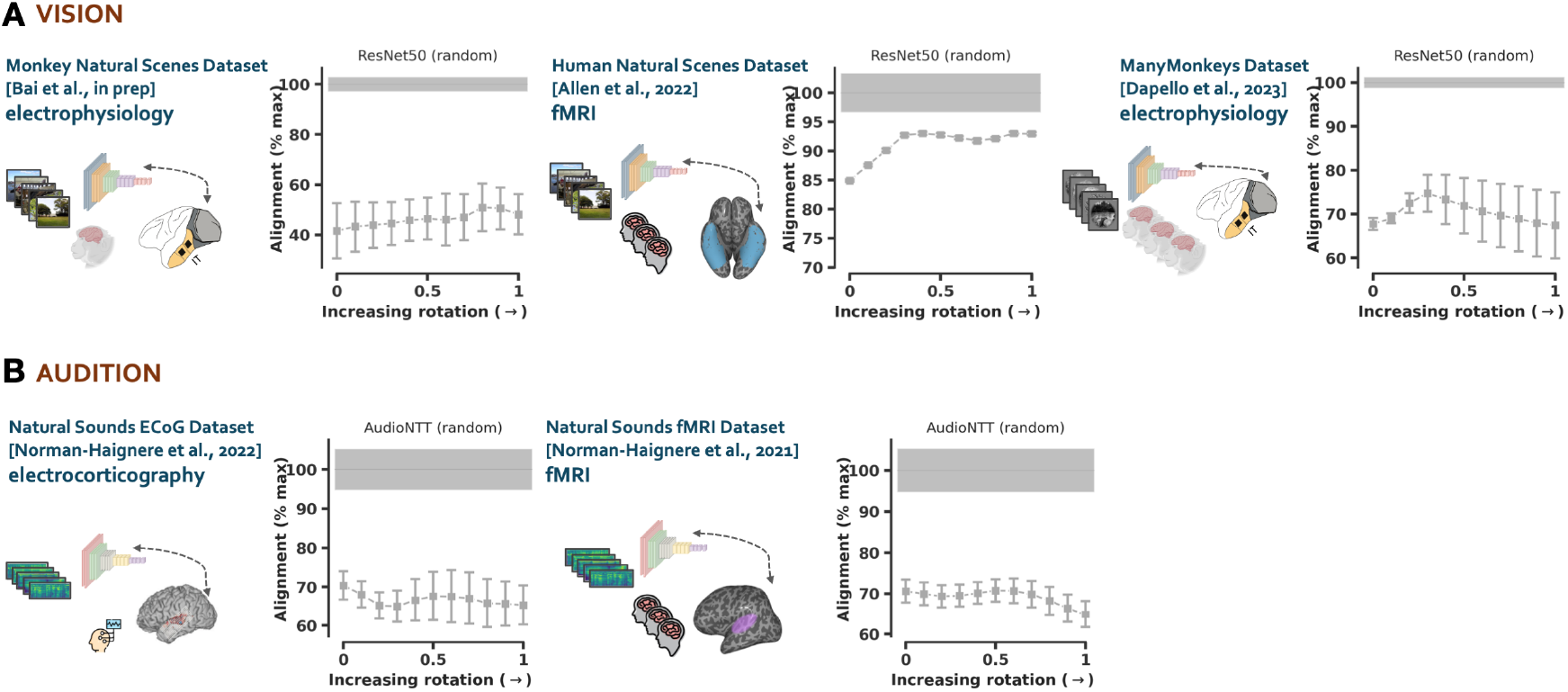
Model-to-brain alignment in untrained networks. (**A**) Alignment of various untrained visual deep convolutional neural networks with biological neural representations in the macaque IT and human ventral visual stream. This alignment is measured using the pairwise matching score and examined as a function of axis rotation (*α*) where *α* = 0 indicates no rotation and *α* = 1 indicates full rotation using a rotation matrix drawn uniformly from SO(n). Alignment is expressed as a proportion of the maximum possible value obtainable across all axis rotations, determined using the ‘Optimal Axis Alignment’ procedure described in the text.(**B**) Alignment of untrained auditory DCNNs with neural representations in the human auditory cortex, again assessed as a function of DCNN axis rotation and measured using the pairwise matching score. Error bars are SEM over multiple splits of the data.

**Fig. S9.**
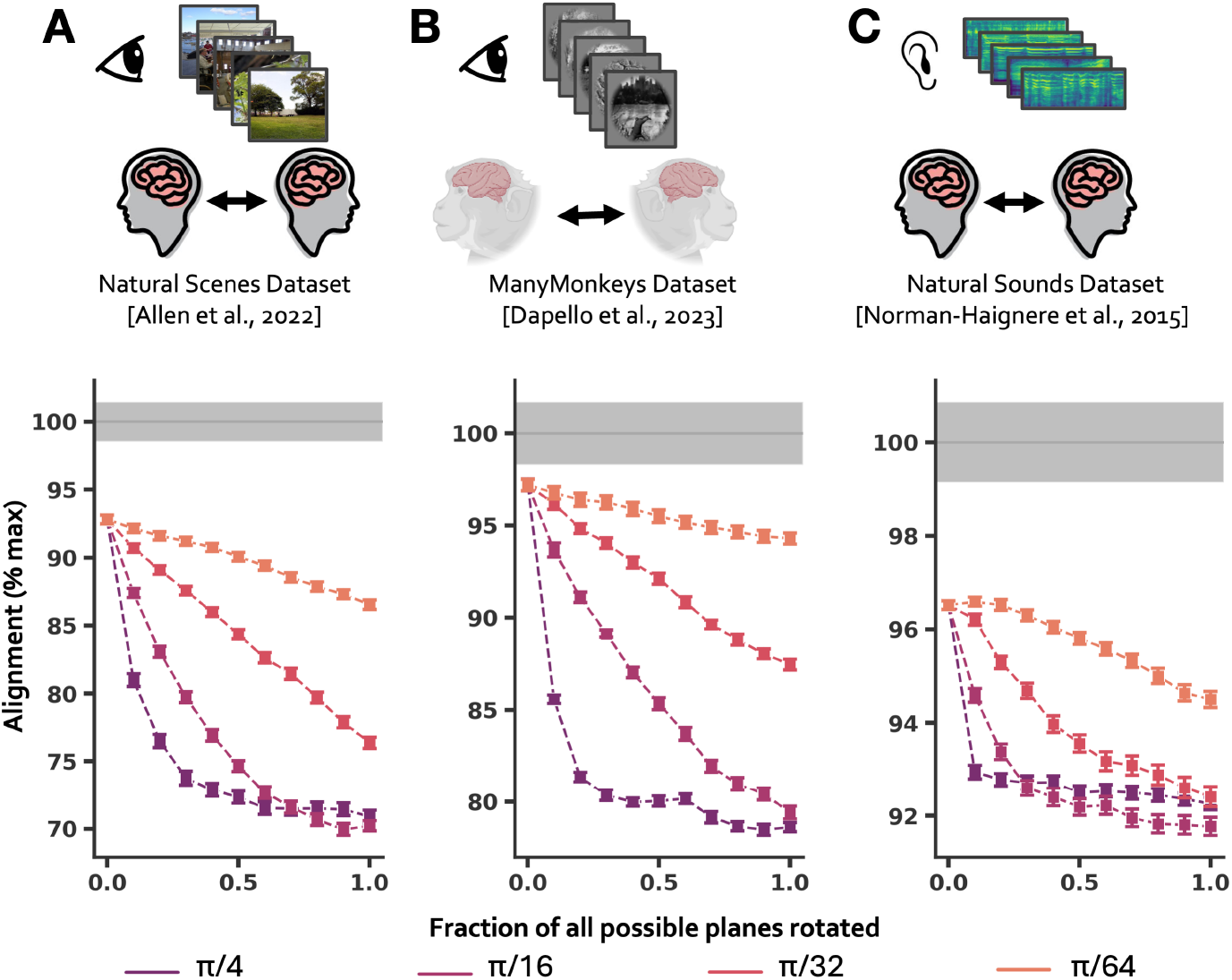
Brain-to-brain alignment under smaller axis perturbations. (**A**), (**B**) and (**C**) depict alignment between conspecifics neural responses in human ventral visual cortex, macaque IT and human auditory cortex, measured using the pairwise matching score. Alignment is examined as a function of the number of plane (2D) rotations (*c*) normalized by the total number of planes (x-axis) and the angle of each 2D rotation (drawn from *π/*4*, π/*16*, π/*32*, π/*64). Error bars are SEM over multiple splits of the data. The results show that higher angles of plane rotation and a greater number of plane rotations more strongly reduce alignment.

**Fig. S10.**
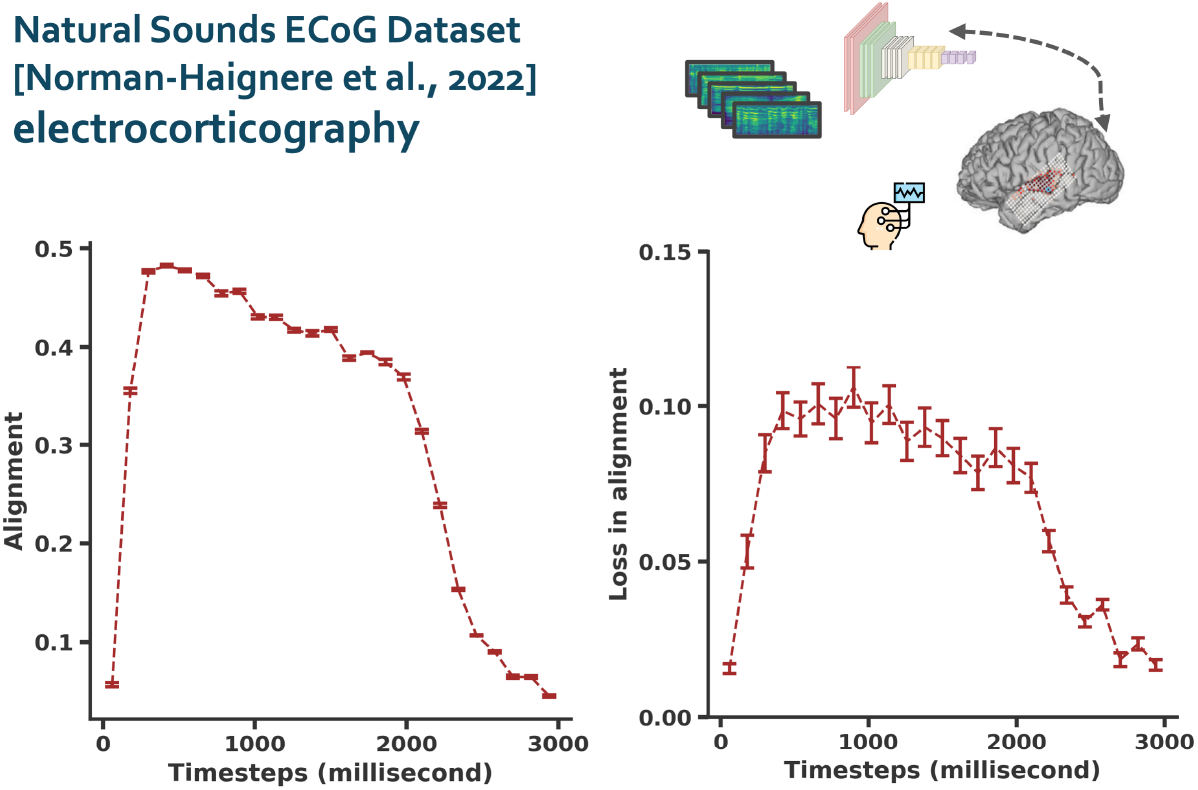
Model-to-brain alignment as a function of time in the auditory domain. Left: Alignment of auditory deep convolutional neural network representations (self-supervised network with AudioNTT architecture) with biological neural representations in the auditory cortex (ECoG recordings) aggregated across different temporal windows following stimulus onset. Alignment is measured using the pairwise matching correlation score. Right: Difference in alignment of the auditory DCNN representation with neural representations in the human auditory cortex aggregated across different temporal windows at no rotation (*α* = 0) and full rotation of the DCNN axis (*α* = 1). Error bars are SEM over multiple splits of the data.

**Fig. S11.**
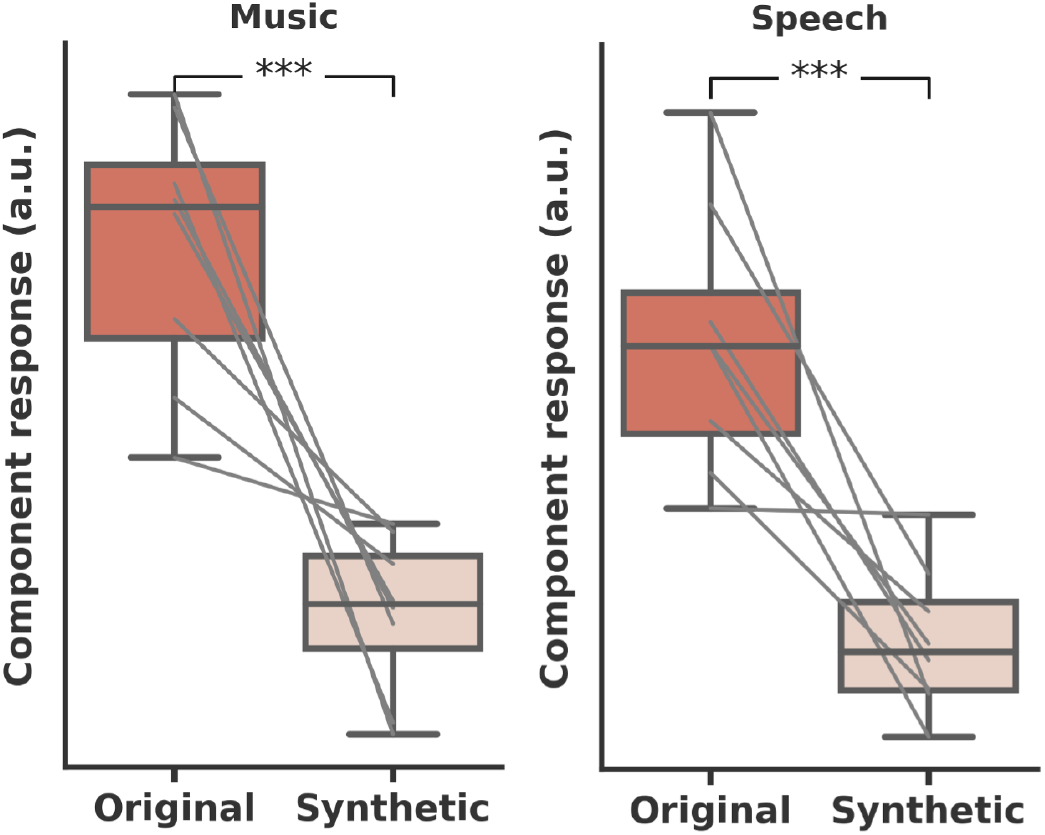
Responses of the music and speech-selective sparse components in the auditory self-supervised DCNN, as obtained by the NMF procedure, to a distinct set of 36 natural speech/music sounds and their corresponding modulation-matched controls. Signed t-test is used to evaluate whether the response to natural sounds was consistently greater than responses to corresponding modulation-matched sounds for these two categories. The modulation-matched synthetic sounds were generated to match the spectrotemporal modulation statistics of the corresponding natural music and speech sounds [105].

**Fig. S12.**
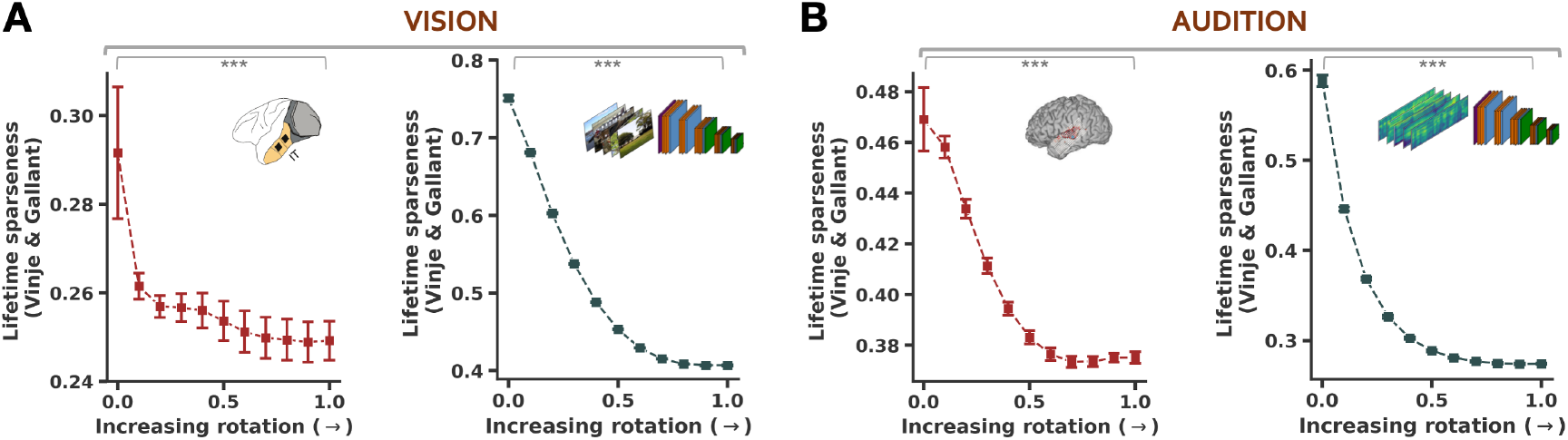
Lineplots show the mean lifetime sparseness, computed using the sparseness measure proposed in Vinje and Gallant (2000), across all units in the representation at different axis rotations. Values near 0 indicate low sparsity, and values near 1 indicate high sparsity. From left to right, the representations include the macaque IT neural responses (from the Macaque-NSD dataset), self-supervised visual DCNN activations, human auditory cortex responses (ECoG) and self-supervised auditory DCNN activations.

**Fig. S13.**
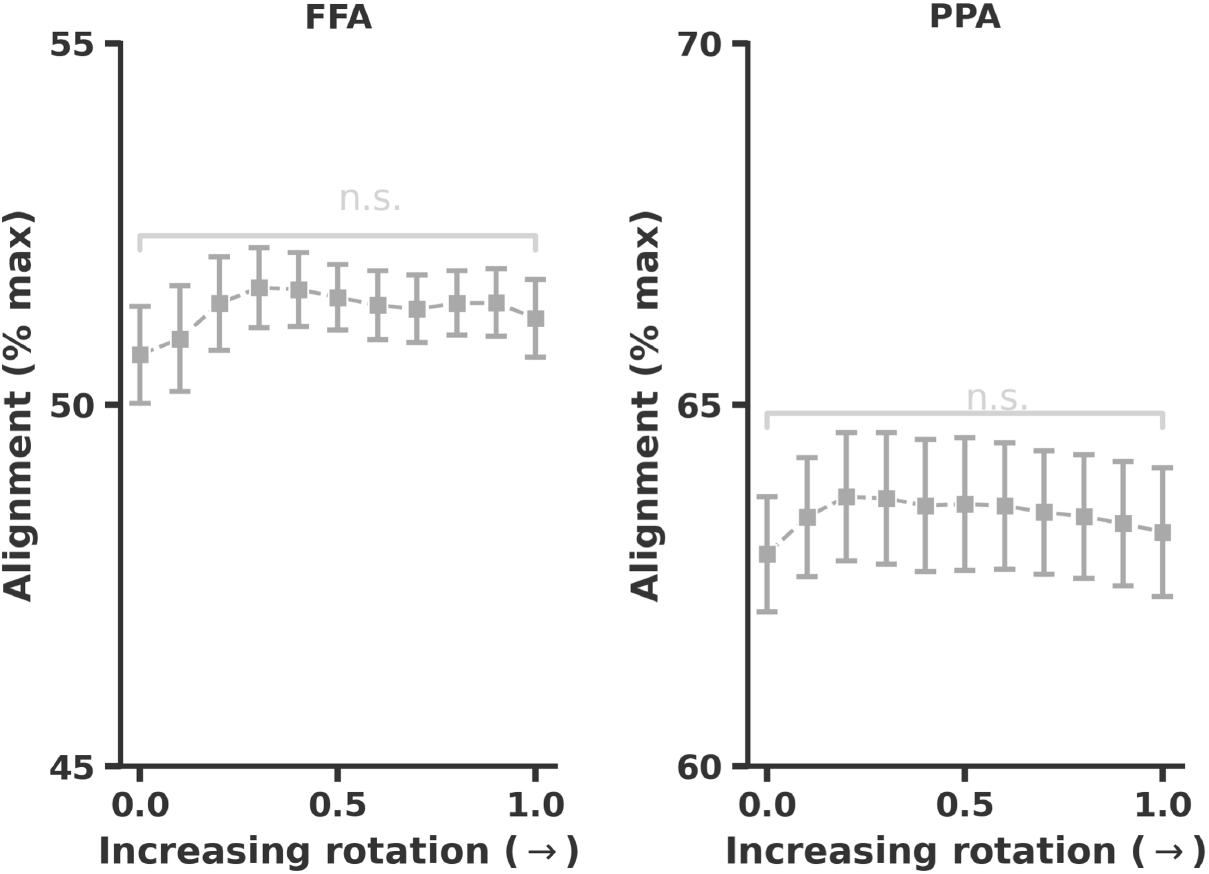
Alignment between conspecifics neural responses in human fusiform face area (FFA, left panel) and the parahippocampal place area (PPA, right panel) as a function of axis rotation, measured using the pairwise matching score. Alignment computed from fMRI activations to the 1000-image NSD stimulus set. Error bars are SEM over multiple splits of the data. Significance tests compare the alignment values at no rotation (*α* = 0) with those at full rotation (*α* = 1). In each subject, the FFA and PPA were defined as the top 100 most face-selective and place-selective voxels, respectively, based on functional localizer experiments and anatomical boundaries provided with NSD. Alignment is expressed as a proportion of the maximal value across all possible axis rotations, assessed using the maximal basis alignment procedure.

